# Structure and function of stator units of the bacterial flagellar motor

**DOI:** 10.1101/2020.05.15.096610

**Authors:** Mònica Santiveri, Aritz Roa-Eguiara, Caroline Kühne, Navish Wadhwa, Howard C. Berg, Marc Erhardt, Nicholas M. I. Taylor

**Affiliations:** Structural Biology of Molecular Machines Group, Protein Structure & Function Program, Novo Nordisk Foundation Center for Protein Research, Faculty of Health and Medical Sciences, University of Copenhagen, Copenhagen, Denmark; Institut für Biologie/Bakterienphysiologie, Humboldt-Universität zu Berlin, Berlin, Germany.; Department of Molecular and Cellular Biology, Harvard University, Cambridge, MA 02138; Rowland Institute at Harvard, Harvard University, Cambridge, MA 02142

## Abstract

Many bacteria use the flagellum for locomotion and chemotaxis. Its bi-directional rotation is driven by the membrane-embedded motor, which uses energy from the transmembrane ion gradient to generate torque at the interface between stator units and rotor. The structural organization of the stator unit (MotAB), its conformational changes upon ion transport and how these changes power rotation of the flagellum, remain unknown. Here we present ~3 Å-resolution cryo-electron microscopy reconstructions of the stator unit in different functional states. We show that the stator unit consists of a dimer of MotB surrounded by a pentamer of MotA. Combining structural data with mutagenesis and functional studies, we identify key residues involved in torque generation and present a mechanistic model for motor function and switching of rotational direction.

**One Sentence Summary:** Structural basis of torque generation in the bidirectional bacterial flagellar motor

## Main Text

Numerous bacteria use rotating flagella to propel themselves (*1, 2*). The ability to move is crucial for bacterial survival and pathogenicity (*3, 4*). The flagellum is made of a long external filament functioning as a propeller; a flexible linking structure, the hook; and a motor embedded in the cell envelope (*5–8*) (Fig. 1A). The ion-powered rotary motor consists of a rotor surrounded by a ring of stator protein complexes (MotAB) that power its rotation (*9–12*). The motor is bidirectional: chemotactic signaling can cause a conformational change in the rotor, known as “switching” (*13*), which results in a change of the rotational direction of the motor.

**Fig. 1.**
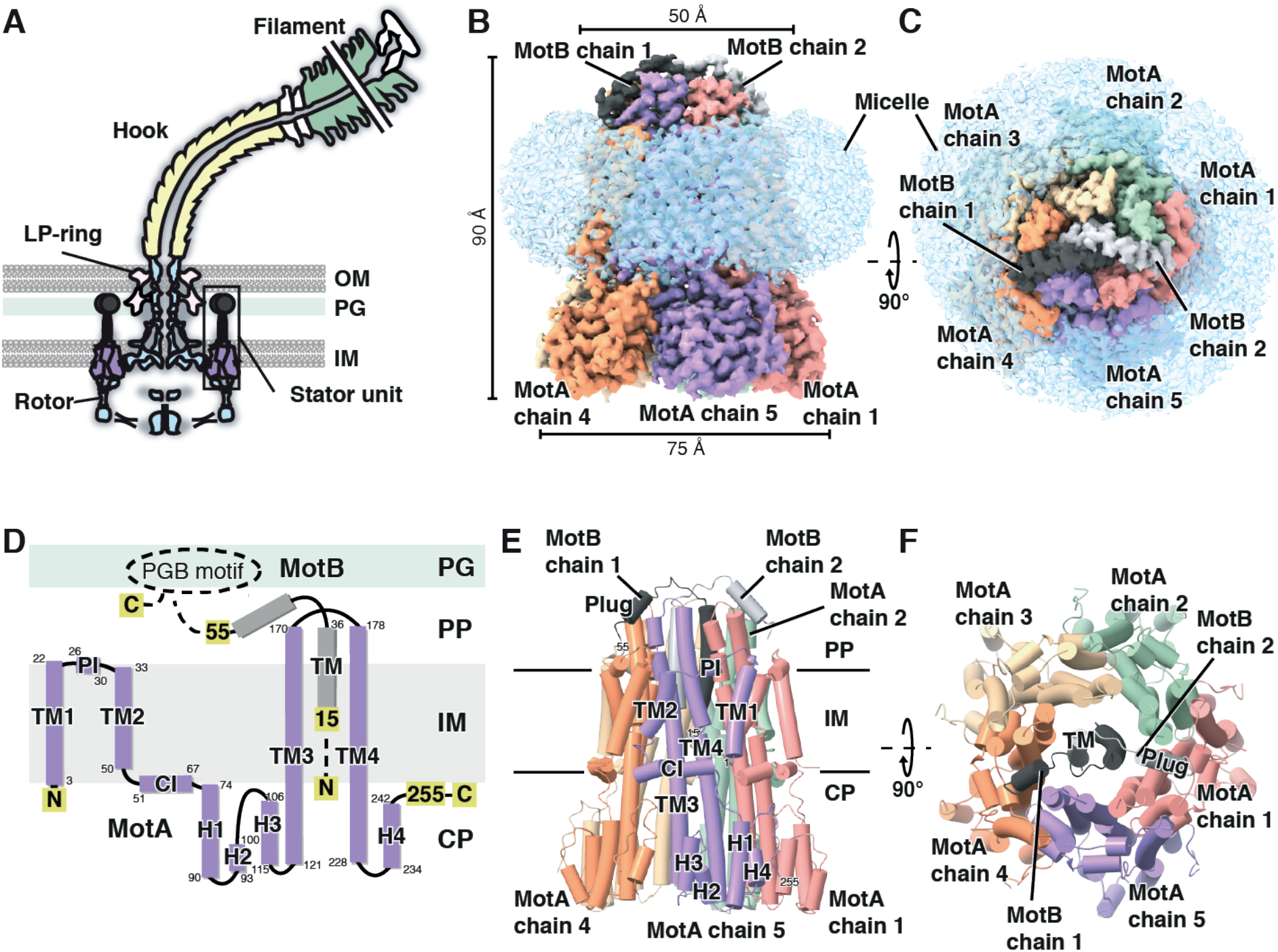
Architecture and topology of the flagellar stator unit MotAB. (**A**) Organization of the bacterial flagellar motor (in gram-negative bacteria). MotA: purple, MotB: dark grey, rotor with export apparatus: light blue, LP-ring: pink, hook: pale yellow, filament: green. Adapted from reference (*62*). OM, outer membrane; PG, peptidoglycan; IM, inner membrane. (**B** and **C**) Side (B) and top (C, periplasmic side) views of the cryo-EM map of the *Cj*MotAB stator unit in a detergent micelle. (**D**) Topology organization of MotA (purple) and MotB (grey) subunit. Dashed lines indicate regions not resolved in this study. The OmpA-like domain containing the PGB motif is indicated as an ellipse. TM helices are numbered from TM1 to TM4. Interface helices are PI for the Periplasmic Interface helix and CI for the Cytosolic Interface helix. Cytosolic helices are numbered from H1 to H5. PG, peptidoglycan; IM, inner membrane; PP, periplasm; CP, cytoplasm; PGB motif, peptidoglycan-binding motif. (**E** and **F**) Side (E) and top (F) views of the atomic model representation. Subunit color code is the same as in (B and C). Secondary structure elements are labelled for MotA chain 5 and MotB chain 1 in (E) and for MotB chain 2 in (F).

Of note, the prokaryotic rotary motor stator unit family (*14*) of which MotAB is the best studied example, is one of only two known motors that use energy from the transmembrane (TM) ion gradient instead of ATP to generate mechanical work (apart from the rotary ATPase family) (*15*). Unlike the rotary ATPases, for which great structural insight has been obtained in recent years (*16*), the mechanism of action of MotAB and stator units of other prokaryotic rotary motors remains poorly understood.

The stator units of the bacterial flagellar motor are embedded in the inner membrane, allowing interaction with the motor and the formation of an ion channel (*9–12, 17, 18*). They are in a plugged, inactive state and get activated upon motor incorporation and peptidoglycan binding (*19*). Rotation of the rotor is powered by dispersion of an ion (generally H^+^ or Na^+^) motive force through the stator units (*20, 21*). It has been proposed that ion binding by the stator unit induces a conformational change in the stator unit itself (*22*). The stator unit protein MotA is thought to contact the FliG protein (through the torque helix (Helix_Torque_) of the C-terminal domain FliGCC) which forms part of the cytoplasmic C-ring of the rotor. In this way, the proposed conformational changes in the stator unit are driving rotation of the rotor (*13, 22–24*). A large body of genetic data is available on mutations in the motor that affect movement and are characterized as Mot^−^ (non-motile, i.e. deficient in motor rotation) or Che^−^ (no chemotaxis, which can be caused by a deficiency in switching rotational direction) (*25*). All previously described mutations in the stator unit proteins are Mot^−^ and not Che^−^. This indicates that switching of the rotational direction is caused solely by structural changes in the rotor. The same conformational changes in the stator unit that power rotation of the rotor in the counterclockwise (CCW) direction, must therefore also power rotation in the clockwise (CW) direction. Upon switching, it is thought that FliGCC makes a 180° turn relative to the stator unit, which allows the rotor to turn in the other direction (*24*).

The stator unit is a complex of two membrane proteins, MotA and MotB (for the H^+^-driven motor) (*26*). MotA contains four TM helices and a large cytoplasmic domain that is proposed to interact with the rotor (*27–29*). MotB contains a single TM helix followed by a large periplasmic domain which can bind peptidoglycan (*30, 31*). The MotB TM domain contains a universally conserved aspartate residue (D22 in *Campylobacter jejuni*, D33 in *Salmonella enterica*), which is thought to be directly involved in proton transport (*32*). Directly following the MotB TM domain is a region known as the plug (*19*). Incorporation of the stator unit in the motor is coupled to the unplugging of the stator unit and peptidoglycan domain dimerization, allowing it to bind peptidoglycan. Crosslinking, biochemical, and genetic data for both MotAB and PomAB (a Na^+^-dependent stator unit) have allowed the identification of residues involved in complex formation and function (*26, 33–36*). Based on these experiments, the stoichiometry of the MotAB stator unit has been suggested to be 4:2. However, this is based on the facts that MotB must at least be a dimer and that the MotA:MotB ratio is at least 2:1 (*37*). Negative stain electron microscopy structures of *Vibrio alginolyticus* PomAB (*38*) and A*quifex aeolicus* MotA (*39*) have been reported, but due to the limited resolution these do not provide information on stator unit stoichiometry or mechanism.

MotAB shows some sequence homology to energizing proteins of other systems, which have been proposed to be stator units of prokaryotic rotary motors (*14*) such as ExbBD (*22*), TolQR (*40*) and AglRQS (*41*). The stoichiometry of ExbBD was uncertain and different experiments reported 4:1, 4:2, 5:2 and 6:3 stoichiometries (*42–44*). However, a recent high-resolution structure of ExbBD shows a clear 5:2 stoichiometry (*45*), which is consistent with the existence of ExbB pentamers in the native *Escherichia coli* membrane (*46*).

Despite great advances in the last decades concerning flagellar motor function, we still do not understand the structural and mechanistic basis of ion transport, channel (un)plugging and torque generation. To help answer these questions, we determined ~3 Å cryo-electron microscopy (cryo-EM) structures of MotAB in different states, as well as lower-resolution structures of several other stator units. Our structures demonstrate a 5:2 stoichiometry for the stator unit complex MotAB, which we show is conserved across the MotAB/PomAB family, and reveal the structural basis of the autoinhibitory plugging of non-incorporated stator units. Furthermore, we infer the structural changes upon proton transport that are driving rotor rotation from the structures of different functional states and validate our structural results using extensive mutagenic analysis of the flagellar stator unit complex. Finally, based on our structural and functional results, we provide a detailed model for motor powering and rotational direction switching.

### The flagellar stator unit is a 5:2 complex

To obtain detailed insight into the mechanism of flagellar stator unit function, we tested the expression and purification of eight H^+^- and Na^+^-dependent stator units of seven different organisms that have been shown to be functional in the presence of C-terminal tags on MotB (figs. S1 and S2). Of the eight protein complexes, six could be purified after detergent solubilization.

For three of these, we obtained cryo-EM reconstructions, with the best resolution (3.1 Å) for *C. jejuni* MotAB (*Cj*MotAB) (fig. S3 and table S1). The maps obtained for *Cj*MotAB allowed building of an atomic model for the nearly complete MotA protein and for the TM helix and plug of MotB (Fig. 1, B to F, and figs. S4 to S7). Therefore, *Cj*MotAB was used as a model system to investigate the structural mechanism of the stator unit. We validated our structures using prior crosslinking, mutational, and tryptophan scanning data of the *E. coli* stator unit (fig. S8, A to F). Structure determination of *Shewanella oneidensis* MotAB (*So*MotAB) and *V. alginolyticus* PomAB (*Va*PomAB) was complicated by preferential orientation of the protein in the ice (figs. S9 and S10), but still allowed clear stoichiometry determination (fig. S3, C to K). We found that *Cj*MotAB forms a 5:2 complex, as do *So*MotAB and *Va*PomAB, suggesting that MotAB stoichiometry is conserved across all flagellar stator units. Furthermore, given the fact that the stoichiometry of ExbBD is identical (*45*), it is likely to be a property of the whole family of stator units of prokaryotic rotary motors.

### Stator unit architecture

The flagellar stator unit has a truncated cone shape (widest at its cytoplasmic region) (Fig. 1B and movie S1). Five copies of MotA cradle the single TM helices of the two copies of MotB. MotA TM helices 3 and 4 make direct interactions with MotB, and both these helices, which span the complete height of MotA, extend to the cytoplasmic domain.

The N-terminal part of MotA forms a parallelogram-like structure in the membrane. It consists of TM helix 1 (crossing from cytoplasm to periplasm), a linker including a 3_10_ helix lying horizontally at the periplasmic side of the membrane (periplasmic interface helix), TM helix 2, crossing from periplasm to cytoplasm and finally the cytoplasmic interface helix, which then connects to the large cytoplasmic domain. Both horizontal helices are very polar on their external sides, and the cytoplasmic interface helix is very basic at its cytoplasmic side (fig. S11, A to D).

The cytoplasmic domain of MotA is made up of two stretches of amino acid chains (residues 69-142 and 211-258) (Fig. 1D). The surface conservation is generally low with two clear exceptions: the MotB interface and a highly conserved region at the bottom of MotA, which extends slightly to the left bottom part of the side of MotA (Fig. 1, D to F, and fig. S11, E to I). The latter region contains residues that have previously been shown to be important for torque generation (*Cj*MotB R89 and E97, corresponding to R90 and E98 in *E. coli*/*S. enterica*) (*47*). Chromosomal point mutants of *S. enterica* MotA (*Se*MotA) R90 and E98 displayed a pronounced defect in motility when the charge of these residues was inverted or the arginine residue was mutated to a smaller amino acid (alanine) (Fig. 2, A and B, and fig. S12). The chromosomal point mutations of MotAB did not affect bacterial growth, suggesting that the observed motility defect was due to impaired motor function and not due to a general deficiency in cellular physiology e.g. increased proton leakage (figs. S13 to S15). In support, charge reversal substitutions in these residues complement charge reversal mutants of oppositely charged residues in the FliG Helix_Torque_ in in *E. coli* (*23*). Therefore, we propose that this part of the structure contacts the rotor, and most likely FliG and its Helix_Torque_, during torque generation (fig. S8, G and H).

**Fig. 2.**
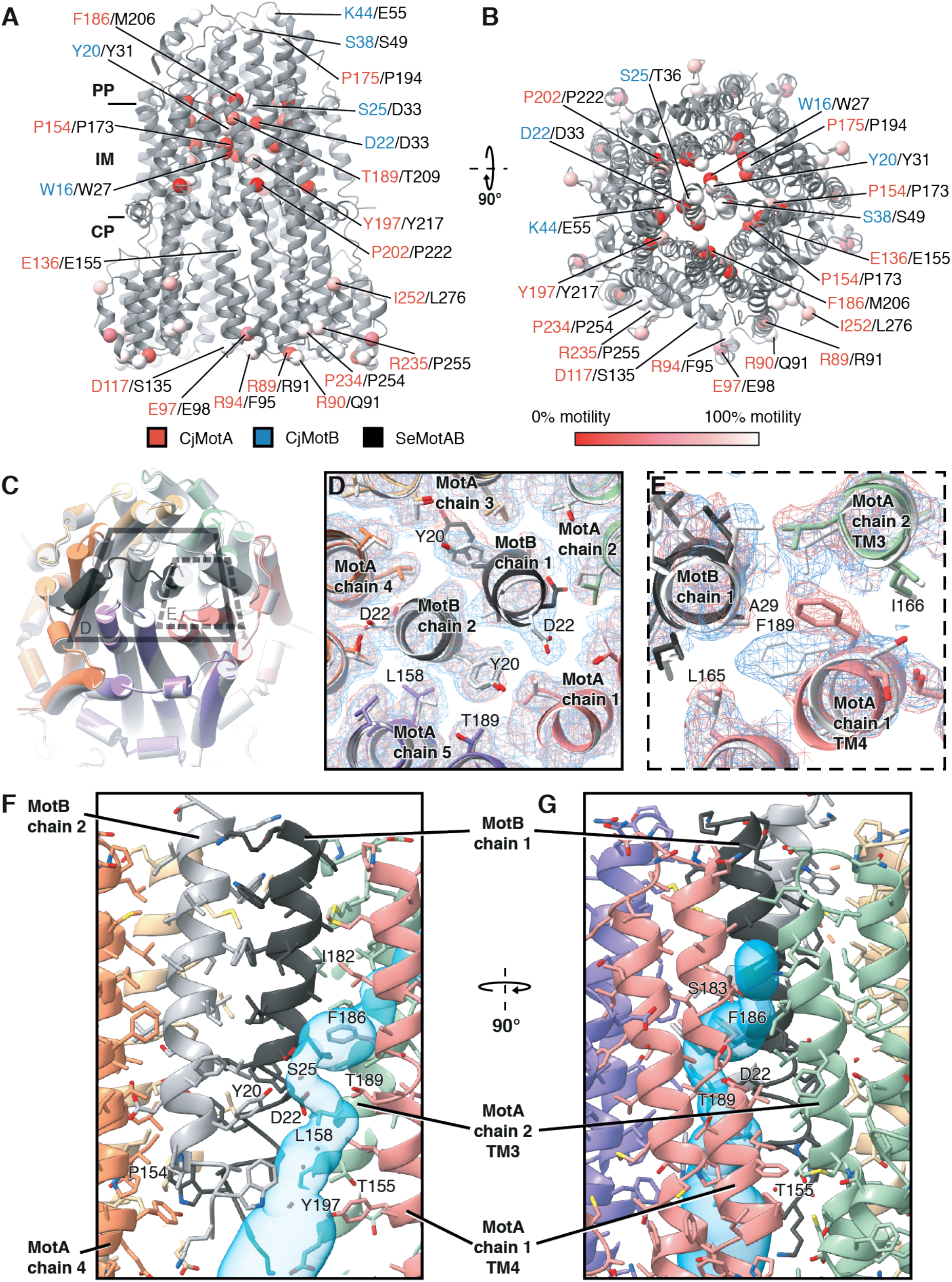
Mutational analysis and conformational changes of the stator unit upon unplugging. (**A** and **B**) Swimming efficiency of *S. enterica* alanine mutants, plotting the mutated residues as spheres on the *Cj*MotAB structure (grey) on the position of the C_α_ atoms of homologous residues in *C. jejuni*. The corresponding *C. jejuni* residue is listed first and colored red (*Cj*MotA) or blue (*Cj*MotB), the residue number of *Se*MotA or *Se*MotB that was mutated is shown in black. (**C**) Superposition of the plugged (colored, same color code as Fig. 1) and unplugged (light grey) models on the periplasmic and TM region of the *Cj*MotAB complex. (**D** and **E**) Close-up view from the periplasmic side of the unplugging effect on the TM plane at the D22 residue level of *Cj*MotB dimer (D) and at the MotA F186 residue (E). The density of the plugged and unplugged stator unit is shown in red and blue, respectively. (**F** and **G**) Side (F) and front (G) views from within the membrane of the predicted channel for the unplugged conformation. A predicted solvent channel accessible to protons and hydronium ions calculated with Mole 2.5 (*63*) (see Methods) is shown in cyan.

The inside of the MotA cytoplasmic region is extremely acidic (fig. S11, A to D). Possibly, this region might act as a reservoir for taking up charges that have passed through the stator unit and/or might interact with the N-terminal tail of MotB, which is visible in our maps but is less ordered than the MotB TM helix (fig. S11, J to L), and which contains various basic residues (fig. S1).

### Stator unit channel (un)plugging

To reveal the active state of the stator unit and the structural basis of unplugging, we determined the 3.0 Å structure of unplugged stator unit *Cj*MotAB(Δ41-60) (which has a deletion of the 20 equivalent MotB residues shown to be important for plugging in *E. coli* (*19*)) and compared it to the full-length *Cj*MotAB structure (Fig. 2, C to E, figs. S5 and S16 and movie S2).

In the full-length structure, MotA interacts with extensions (or plugs, one per MotB chain) immediately C-terminal of the MotB TM helix. Seen from the periplasmic side of the channel, the plugs have pseudo-mirror symmetry, resulting in extensive interaction between both plugs at the crossover point (Fig. 2C). After a short coil structure (residues 40-44), both plugs form a helix which lies in between MotA subunits, with three MotA subunits on one side and two on the other. Deletion of the plug region in *E. coli* and *S. enterica* MotB results in a massive influx of protons into the cytoplasm and inhibition of cell growth (*19, 48*), therefore the plug region is important to prevent proton leakage.

Comparing the structures of plugged and unplugged stator units, few conformational changes can be observed based on the lowest C_α_ root-mean-square deviation (r.m.s.d.) superposition (0.714 Å) (Fig. 2C). For the larger residues of *Cj*MotB, changes are limited to different conformations of Y20, D22 and F23 in chain 1. Looking at the universally conserved *Cj*MotB D22 residue, we note that one (*Cj*MotB chain 1 D22) is mostly accessible (but pointing away) from the cytoplasmic interface (where we can observe solvent molecules), whereas the other (*Cj*MotB chain 2 D22) is interacting with MotA and not accessible to solvent, both in the plugged and unplugged structures (Fig. 2D). This suggests that *Cj*MotB chain 1 D22, but not *Cj*MotB chain 2 D22, would be protonatable and/or able to interact with hydronium.

The MotB TM helix and the internal MotB-interacting surface of MotA are highly conserved (fig. S11, E to I). Their interaction surfaces are almost purely hydrophobic, the only polar or charged residues are *Cj*MotA T155 and T189 and *Cj*MotB Y20, D22 and S25 (Fig. 2, F and G). The corresponding *Se*MotAB residues are also the only polar residues in this region (fig. S17). Of these, *Cj*MotA T189 and *Cj*MotB D22 are universally conserved and the polarity of *Cj*MotB S25 (which can be threonine in some stator units) is conserved as well (fig. S1). Interestingly, all these residues lie at the height of the bottom part of the MotB TM helix, or put differently, at or below the height of the inner membrane. Using swimming motility assays, we show that in *S. enterica*, only *Se*MotB D33 (*Cj*MotB D22) is absolutely required for motor function, but *Se*MotA T209A (*Cj*MotA T189) also displays severely decreased motility (while not affecting growth) (Fig. 2, A and B, figs. S12 to S15 and table S2).

These observations suggest that an access pathway must exist for protons and/or hydronium ions to MotB chain 1 D22 from the periplasm in the unplugged structure, but not in the plugged structure. Such a pathway appears to exist from the side of MotA between chains 1 and 2, just above the TM region. In the unplugged structure, but not in the plugged structure, MotA chain 1 F186 is present in two alternate positions (positions 1 and 2), as can be clearly seen in the map (Fig. 2E and fig. S18). Position 1 is the same as in the plugged structure. Position 2, which appears to be the most occupied, overlaps with the location that in the plugged structure is taken up by a solvent molecule. This position is also in close proximity to MotB chain 1 S25 (and a solvent molecule that can be found near this residue in both structures), D22 and Y20 and MotA chain 1 T189. The polar residues outlined before appear to form a solvent-accessible channel (Fig. 2, F and G). The channel is lined with residues that have previously been shown to be important for ion transport (*49–51*) and/or are differentially conserved between H^+^- and Na^+^-dependent stator units (figs. S1 and S19). *Cj*MotA F186 (*Se*MotA M206) is universally conserved hydrophobic residue (fig. S1). We found that mutations of *Se*MotA M206 to a small amino acid (M206A) or negatively charged amino acid (M206D) completely abrogated motility while not affecting growth (Fig. 2, A and B, figs. S12 to S15 and table S2). This supports previous findings that M206 is involved in torque generation and proton translocation as well as pH-dependent stator assembly (*52*). We conclude that *Cj*MotA F186 is a hydrophobic residue shielding the periplasm from the hydrophilic channel of MotAB. We propose that unplugging increases flexibility of *Cj*MotA F186, allowing the passage of protons or hydronium ions through the channel.

### Conformational changes upon proton transport

To gain insight into the conformational changes that *Cj*MotAB undergoes upon proton transport, we determined the 3.0 Å cryo-EM structure of stator units that combine the unplugging mutation *Cj*MotB(Δ41-60) with a *Cj*MotB(D22N) mutation, mimicking protonation or hydronium binding of D22 (Fig. 3, A and B, figs. S6 and S16 and movie S3). The structure of *Cj*MotAB(Δ41-60, D22N) is extremely similar to *Cj*MotAB(Δ41-60) when observing the lowest Cα r.m.s.d. (0.297 Å) superposition, with one exception in MotB chain 1 (nomenclature based on structure alignment with lowest Cα r.m.s.d. not taking into account large-scale rotational movement). N22 is clearly in a different position compared to D22, pointing down towards the cytoplasmic interface, where we can also distinguish several putative solvent molecules (fig. S11, M to O). We conclude that proton or hydronium binding or release by *Cj*MotB chain 1 D22 establishes a small conformational change in and around this residue, strongly suggesting that this residue is directly involved in the shuttling of protons or hydronium ions.

**Fig. 3.**
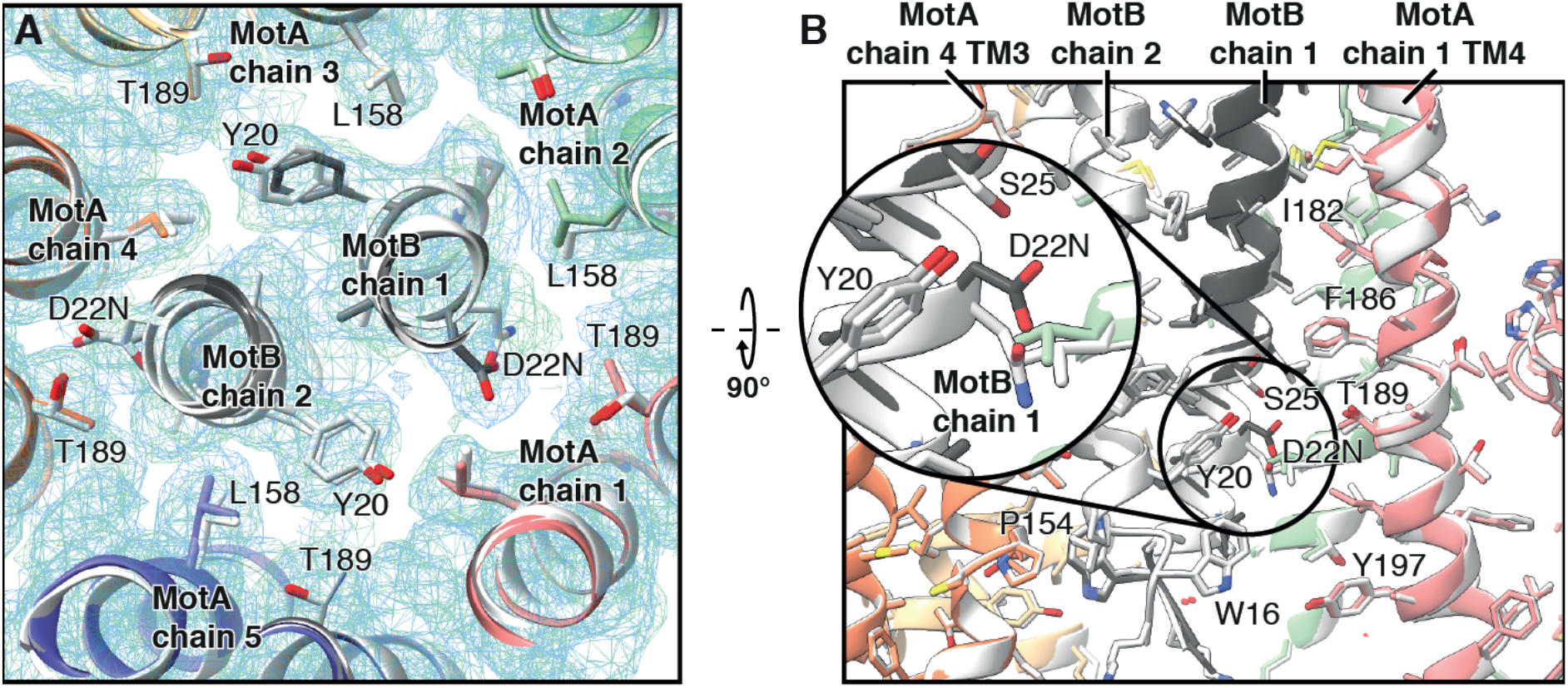
Conformational changes upon (mimicking of) (de)protonation. (**A**) Superposition of *Cj*MotAB(Δ41-60) (light grey) and *Cj*MotAB(Δ41-60, D22N) (same color code as Fig. 1) in a close-up view from the periplasmic side. The proton- or hydronium-bound state is mimicked by the mutation D22N. The density maps are shown for both, *Cj*MotAB(Δ41-60) (blue) and *Cj*MotAB(Δ41-60, D22N) (green). (**B**) Same as (A) but a side view from within the membrane. The inset shows a magnification of the region around *Cj*MotB chain 1 residue D22/N22, illustrating the conformational change around this residue upon mutation.

### A rotational model for torque generation

Stator units power the rotation of the flagellar motor using energy derived from the ion motive force. As mentioned, the only motors harnessing ion motive force to generate work found in nature are the stator unit family of prokaryotic rotary motors (exemplified by MotAB) and rotary ATPases. Our analysis combined with a plethora of prior structural and functional data shows that the stator units interact with the rotor through the cytoplasmic domains of MotA to provide torque. Two mechanisms can be proposed for how torque is generated: a “rotational” model, where MotA rotates around MotB, and a “large conformational change” model, where MotAB changes between two conformations without rotation of MotA around MotB.

Our results are fully consistent with a rotational mechanism of the stator unit, rather than a large conformational change mechanism. The C_α_ r.m.s.d. between *Cj*MotAB(Δ41-60) and *Cj*MotAB(Δ41-60, D22N) is 0.297 Å. It has been estimated that on the order of 37 (*53*) or 70 (*54*) ions are turned over per stator unit, per rotation of the rotor. From the geometry of the motor (fig. S20) we calculate that the rotor needs to traverse an arc length of ~20-38 Å per ion. Consequently, the observed conformational changes are approximately two orders of magnitude smaller than the estimated arc length traversed per ion passage. Rotations of 36° or 72° of MotA around MotB, however, would traverse arc lengths of 24 and 47 Å, respectively. Therefore, we propose that MotAB, and most likely all stator unit proteins of prokaryotic rotary motors, use a rotational mechanism to perform work.

### An inchworm model for powering of rotation of MotA around MotB

Given the structural similarity of both unplugged structures and the number of ions per stator and per rotor rotation, it follows that rotation of MotA around MotB will occur in steps of either 36° (so that after each rotary step, MotB chain 2 would be in a position equivalent, with respect to MotA, to where MotB chain 1 was before the rotation, and *vice versa*) or 72° (with MotB chain 1 and chain 2 in the same equivalent positions, with respect to MotA, which they had before the rotation). The first model (36° rotation) is more likely, as in this model the universally conserved aspartate residue of both chains would transport ions alternately, while this would not happen in the second model. Superposing the *Cj*MotAB(Δ41-60) and *Cj*MotAB(Δ41-60, D22N) structures in this way and making the natural assumption that the hydrophobic MotA interior can only rotate around charge-neutralized MotB D22 readily points to a model for how rotation occurs at the molecular level (fig. S21 and movies S4 and S5). Note that charge neutralization by proton binding of a carboxylate group is also used in the Fo/Vo/Ao component of rotary ATPases, where protonation of an aspartate or glutamate residue on the c protein allows that residue’s entry into the hydrophobic interior of the lipid membrane and therefore rotation of the c-ring (*16, 55*). The protein geometry suggests a clockwise rotation (when observed from the extracellular/periplasmic side) of MotA around MotB: MotB chain 1 N22 is close to the equivalent position taken up by D22/N22 in MotB chain 2 (near T189), located clockwise.

The proposed mechanism is very akin to human-made inchworm motors. Each MotB D22 alternately engages with MotA, in a site between *Cj*MotA T189, P154 and G150. When MotB D22 is engaged, it can (help) drive a power stroke (when the charge of the other MotB D22 becomes neutralized). When it is not engaged, it picks up a proton from the channel and inches to the position where it can (help) drive the power stroke. The mentioned MotA residues at the site of engagement are extensively conserved across all stator units of rotary prokaryote motors (*56*). Furthermore, the *Va*PomA T186A mutation abrogates motility as well as Na^+^-dependent structural changes in *Va*PomAB (*51*) and the corresponding *Se*MotA T209A mutation has a severe motility defect (Fig. 2, A and B, and fig. S12), as mentioned (both corresponding to *Cj*MotA T189).

### Powering bidirectional rotation of the flagellum

In the following, we present a simple but comprehensive model integrating the data presented here, prior data and previous models for stator unit activation, torque generation and directional switching.

Before association with the rotor and peptidoglycan binding, MotAB is in the plugged state and the channel is closed. Association with the rotor and peptidoglycan binding is coupled to unplugging of the channel (Fig. 4A). The cytoplasmic domains of MotAB incorporated in the motor are located such that (at least) one of them can interact with FliG Helix_Torque_ (Fig. 4B). Based on our structural data, genetic data (*23*) and our modeling of the FliG–MotA interaction (fig. S8, G and H), FliG structural data (*24*) and tomographic data on the *Borrelia* flagellar motor (*57*), the rotor in the CCW state interacts with the inside (the side facing the motor axis) of the stator unit. Upon proton or hydronium binding and release by MotB D22, MotA rotates CW, relative to MotB, which in turn moves the rotor in CCW direction, as MotB is stably anchored to the peptidoglycan. Note that CW rotation of MotA is also predicted by our model outlined in the previous section (fig. S21).

**Fig. 4.**
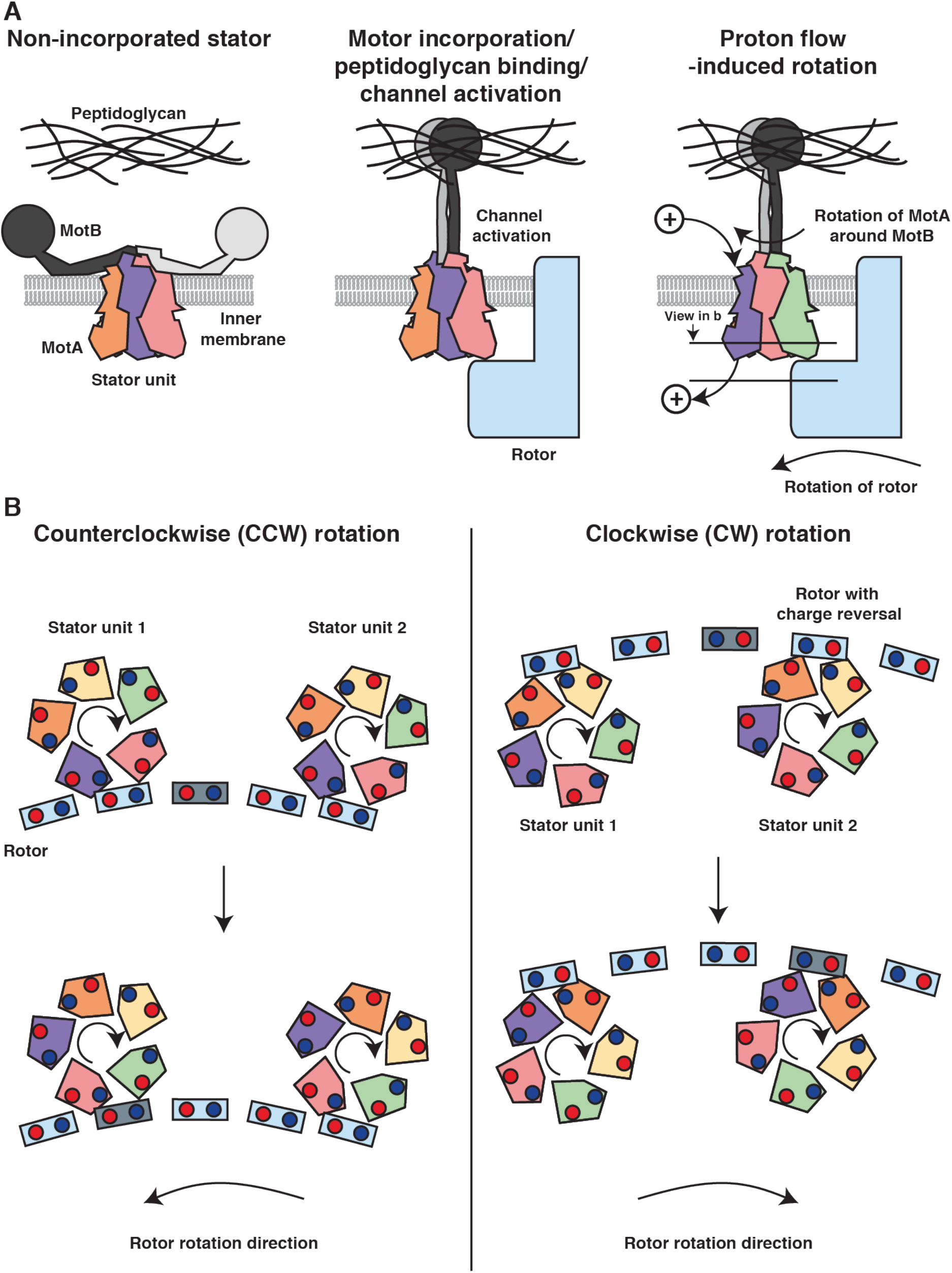
Models of MotAB activation and function. (**A**) Activation mechanism of MotAB. MotB of non-incorporated stator units plugs the proton channel. Motor incorporation is coupled to MotB peptidoglycan binding domain dimerization and peptidoglycan binding. This activates the channel. Proton or hydronium binding and release by the universally conserved MotB aspartate residue (*Cj*MotB D22, *Se*MotB D33) will generate rotation of the MotA pentamer around the MotB dimer, which in turn powers the rotation of the flagellar rotor. MotA and MotB: multi-colored (same color code as Fig. 1). A proton or hydronium is represented by a sphere with a + symbol. (**B**) Torque generation mechanism during default rotation (CCW, left) and after switching direction (CW, right). Two stator units are shown in top view from the flagellum/extracellular side of the motor. FliG Helix_Torque_ of 5 copies of FliG are shown. MotA: same color code as Fig. 1, FliG Helix_Torque_: light blue with 1 copy highlighted in grey blue. Conserved acidic and basic residues (in MotA and FliG Helix_Torque_) are symbolized with red and blue circles, respectively. See fig. S8, G to H, for the modeled MotA–FliG interaction. Rotation directions are given for a motor observed from the extracellular side.

Upon CheY-P-induced directional switching, FliGCC (and therefore FliG Helix_Torque_) makes a ~180° turn relatively to the stator unit (*58*). We propose that the geometry of the stator unit rotor interface allows FliG Helix_Torque_ to now engage the outside (the side facing away from the motor axis) of the MotA pentamer. The rotation of MotA relative to MotB is still the same (CW), but because of the changed positioning of FliG, the same conformational change in MotA now powers rotation of the rotor in the CW direction (Fig. 4B).

The model is consistent with the recently observed structural changes in the C-ring of CCW-biased and CW-locked mutants of *V. alginolyticus* (*59*). Furthermore, it predicts that the reversal of the ion motive force would invert the rotation direction of the stator unit and hence of the motor. This has previously been observed in *E. coli* (*60*), lending experimental support to our model. Furthermore, our model does not require any different conformational changes for the stator unit in powering rotation of the rotor in the CCW vs. CW directions, consistent with the apparent lack of Che^−^ mutations in MotA and MotB. According to our model (fig. S21) and the geometry of the rotor and stator unit (fig. S20), binding and release of two protons (two 36° rotations) allow the stator unit to bind the neighboring FliG molecule, or a total of 68 protons per stator unit per rotation of the C-ring (assuming 34 FliG molecules per C-ring), in good agreement with previous estimates (*54*). The geometry, jointly with the proposed inchworm mechanism of the stator unit, is also consistent with the observed high duty ratio of the motor (*61*), as the implied handover mechanisms allow that rotor and stator as well as MotA and MotB remain firmly associated all the time.

In summary, we provide here fundamental insight into stator unit organization and a biophysical model of torque generation and switching of rotational direction of the flagellar motor. These results provide a structure-based framework for a profusion of experiments on stator units of prokaryote rotatory motors, the bacterial flagellar motor and nanoscale motors in general.

## Acknowledgments

We thank the Danish Cryo-EM Facility at the Core Facility for Integrated Microscopy (CFIM) at the University of Copenhagen and Tillmann Pape for support during data collection. Part of the data processing was performed at the Computerome, the Danish National Computer for Life Sciences. We thank Haidai Hu, Guillermo Montoya and Carlos Fernández Tornero for feedback on the manuscript.

## Author contributions

M.S. cloned, expressed, purified, prepared all cryo-grids, collected cryo-EM data and determined the structure of *Cj*MotAB (wild-type and mutants), *So*MotAB and *Va*PomAB. A.R.E. helped M.S. with protein expression and purification and prepared figures for the manuscript together with N.M.I.T. C.K. generated chromosomal *Se*MotAB mutants, performed and analyzed motility assays and growth curves. C.K. and M.E. interpreted motility and growth curve data and prepared figures for the manuscript. N.M.I.T. planned the experiments, refined and validated the structures and wrote the first draft of the paper, which was then edited by M.E., and analyzed by N.W. and H.C.B. All authors contributed to the revision of the manuscript.

## Data and materials availability

Atomic coordinates for *Cj*MotAB, *Cj*MotAB(Δ41-60), and *Cj*MotAB(Δ41-60, D22N) were deposited in the Protein Data Bank under accession codes 6YKM, 6YKP and 6YKR. The corresponding electrostatic potential maps were deposited in the Electron Microscopy Data Bank (EMDB) under accession codes EMD-10828, EMD-10829 and EMD-10830, respectively. The electrostatic potential maps for *So*MotAB and *Va*PomAB were deposited in the EMDB under accession codes EMD-10831 and EMD-10832, respectively. All other data are available from the corresponding author upon reasonable request

## Supplementary Materials

Materials and Methods Figures S1 – S21

Tables S1 – S2

Movies S1 – S5

References (64 – 88)

## References and Notes

1. M. Silverman, M. Simon, Flagellar rotation and the mechanism of bacterial motility. Nature 249, 73–74 (1974).

2. H. C. Berg, R. A. Anderson, Bacteria swim by rotating their flagellar filaments. Nature 245, 380–382 (1973).

3. Q. Duan, M. Zhou, L. Zhu, G. Zhu, Flagella and bacterial pathogenicity. Journal of basic microbiology 53, 1–8 (2013).

4. J. Haiko, B. Westerlund-Wikström, The role of the bacterial flagellum in adhesion and virulence. Biology 2, 1242–1267 (2013).

5. S. Nakamura, T. Minamino, Flagella-Driven Motility of Bacteria. Biomolecules 9, 279 (2019).

6. H. C. Berg, The rotary motor of bacterial flagella. Annual review of biochemistry 72, 19–54 (2003).

7. D. J. DeRosier, The turn of the screw: the bacterial flagellar motor. Cell 93, 17–20 (1998).

8. Y. V. Morimoto, T. Minamino, Structure and function of the bi-directional bacterial flagellar motor. Biomolecules 4, 217–234 (2014).

9. J. W. Coulton, R. G. Murray, Cell envelope associations of Aquaspirillum serpens flagella. Journal of bacteriology 136, 1037–1049 (1978).

10. S. Khan, M. Dapice, T. S. Reese, Effects of mot gene expression on the structure of the flagellar motor. Journal of molecular biology 202, 575–584 (1988).

11. S. Khan, I. H. Khan, T. S. Reese, New structural features of the flagellar base in Salmonella typhimurium revealed by rapid-freeze electron microscopy. Journal of bacteriology 173, 2888–2896 (1991).

12. S. Khan, D. M. Ivey, T. A. Krulwich, Membrane ultrastructure of alkaliphilic Bacillus species studied by rapid-freeze electron microscopy. Journal of bacteriology 174, 5123–5126 (1992).

13. T. Minamino, M. Kinoshita, K. Namba, Directional Switching Mechanism of the Bacterial Flagellar Motor. Computational and structural biotechnology journal 17, 1075–1081 (2019).

14. Y.-W. Lai, P. Ridone, G. Peralta, M. M. Tanaka, M. A. B. Baker, Evolution of the stator elements of rotary prokaryote motors. Journal of bacteriology, (2019).

15. K. K. Mandadapu, J. A. Nirody, R. M. Berry, G. Oster, Mechanics of torque generation in the bacterial flagellar motor. Proceedings of the National Academy of Sciences of the United States of America 112, E4381–4389 (2015).

16. W. Kühlbrandt, K. M. Davies, Rotary ATPases: A New Twist to an Ancient Machine. Trends in biochemical sciences 41, 106–116 (2016).

17. J. Stader, P. Matsumura, D. Vacante, G. E. Dean, R. M. Macnab, Nucleotide sequence of the Escherichia coli motB gene and site-limited incorporation of its product into the cytoplasmic membrane. Journal of bacteriology 166, 244–252 (1986).

18. M. L. Wilson, R. M. Macnab, Overproduction of the MotA protein of Escherichia coli and estimation of its wild-type level. Journal of bacteriology 170, 588–597 (1988).

19. E. R. Hosking, C. Vogt, E. P. Bakker, M. D. Manson, The Escherichia coli MotAB proton channel unplugged. Journal of molecular biology 364, 921–937 (2006).

20. S. H. Larsen, J. Adler, J. J. Gargus, R. W. Hogg, Chemomechanical coupling without ATP: the source of energy for motility and chemotaxis in bacteria. Proceedings of the National Academy of Sciences of the United States of America 71, 1239–1243 (1974).

21. N. Hirota, Y. Imae, Na+-driven flagellar motors of an alkalophilic Bacillus strain YN-1. The Journal of biological chemistry 258, 10577–10581 (1983).

22. S. Kojima, D. F. Blair, Conformational change in the stator of the bacterial flagellar motor. Biochemistry 40, 13041–13050 (2001).

23. J. Zhou, S. A. Lloyd, D. F. Blair, Electrostatic interactions between rotor and stator in the bacterial flagellar motor. Proceedings of the National Academy of Sciences of the United States of America 95, 6436–6441 (1998).

24. L. K. Lee, M. A. Ginsburg, C. Crovace, M. Donohoe, D. Stock, Structure of the torque ring of the flagellar motor and the molecular basis for rotational switching. Nature 466, 996–1000 (2010).

25. S. Yamaguchi, H. Fujita, A. Ishihara, S. Aizawa, R. M. Macnab, Subdivision of flagellar genes of Salmonella typhimurium into regions responsible for assembly, rotation, and switching. Journal of bacteriology 166, 187–193 (1986).

26. H. Tang, T. F. Braun, D. F. Blair, Motility protein complexes in the bacterial flagellar motor. Journal of molecular biology 261, 209–221 (1996).

27. G. E. Dean, R. M. Macnab, J. Stader, P. Matsumura, C. Burks, Gene sequence and predicted amino acid sequence of the motA protein, a membrane-associated protein required for flagellar rotation in Escherichia coli. Journal of bacteriology 159, 991–999 (1984).

28. D. F. Blair, H. C. Berg, Mutations in the MotA protein of Escherichia coli reveal domains critical for proton conduction. Journal of molecular biology 221, 1433–1442 (1991).

29. J. Zhou, R. T. Fazzio, D. F. Blair, Membrane topology of the MotA protein of Escherichia coli. Journal of molecular biology 251, 237–242 (1995).

30. S. Kojima et al., The Helix Rearrangement in the Periplasmic Domain of the Flagellar Stator B Subunit Activates Peptidoglycan Binding and Ion Influx. Structure (London, England: 1993) 26, 590–598.e595 (2018).

31. A. Roujeinikova, Crystal structure of the cell wall anchor domain of MotB, a stator component of the bacterial flagellar motor: implications for peptidoglycan recognition. Proceedings of the National Academy of Sciences of the United States of America 105, 10348–10353 (2008).

32. J. Zhou et al., Function of protonatable residues in the flagellar motor of Escherichia coli: a critical role for Asp 32 of MotB. Journal of bacteriology 180, 2729–2735 (1998).

33. K. Sato, M. Homma, Multimeric structure of PomA, a component of the Na+-driven polar flagellar motor of vibrio alginolyticus. The Journal of biological chemistry 275, 20223–20228 (2000).

34. T. F. Braun, D. F. Blair, Targeted disulfide cross-linking of the MotB protein of Escherichia coli: evidence for two H(+) channels in the stator Complex. Biochemistry 40, 13051–13059 (2001).

35. L. L. Sharp, J. Zhou, D. F. Blair, Features of MotA proton channel structure revealed by tryptophan-scanning mutagenesis. Proceedings of the National Academy of Sciences of the United States of America 92, 7946–7950 (1995).

36. L. L. Sharp, J. Zhou, D. F. Blair, Tryptophan-scanning mutagenesis of MotB, an integral membrane protein essential for flagellar rotation in Escherichia coli. Biochemistry 34, 9166–9171 (1995).

37. S. Kojima, D. F. Blair, Solubilization and purification of the MotA/MotB complex of Escherichia coli. Biochemistry 43, 26–34 (2004).

38. K. Yonekura, S. Maki-Yonekura, M. Homma, Structure of the Flagellar Motor Protein Complex PomAB: Implications for the Torque-Generating Conformation. Journal of bacteriology 193, 3863–3870 (2011).

39. N. Takekawa et al., The tetrameric MotA complex as the core of the flagellar motor stator from hyperthermophilic bacterium. Scientific reports 6, 31526 (2016).

40. E. Cascales, R. Lloubes, J. N. Sturgis, The TolQ-TolR proteins energize TolA and share homologies with the flagellar motor proteins MotA-MotB. Molecular microbiology 42, 795–807 (2001).

41. M. Sun, M. Wartel, E. Cascales, J. W. Shaevitz, T. Mignot, Motor-driven intracellular transport powers bacterial gliding motility. Proceedings of the National Academy of Sciences of the United States of America 108, 7559–7564 (2011).

42. H. Celia et al., Structural insight into the role of the Ton complex in energy transduction. Nature 538, 60–65 (2016).

43. A. Sverzhinsky et al., Membrane Protein Complex ExbB4-ExbD1-TonB1 from Escherichia coli Demonstrates Conformational Plasticity. Journal of bacteriology 197, 1873–1885 (2015).

44. S. Maki-Yonekura et al., Hexameric and pentameric complexes of the ExbBD energizer in the Ton system. eLife 7, (2018).

45. H. Celia et al., Cryo-EM structure of the bacterial Ton motor subcomplex ExbB-ExbD provides information on structure and stoichiometry. Communications biology 2, 358–356 (2019).

46. D. S. Chorev et al., Protein assemblies ejected directly from native membranes yield complexes for mass spectrometry. Science (New York, NY) 362, 829–834 (2018).

47. J. Zhou, D. F. Blair, Residues of the cytoplasmic domain of MotA essential for torque generation in the bacterial flagellar motor. Journal of molecular biology 273, 428–439 (1997).

48. Y. V. Morimoto, Y.-S. Che, T. Minamino, K. Namba, Proton-conductivity assay of plugged and unplugged MotA/B proton channel by cytoplasmic pHluorin expressed in Salmonella. FEBS letters 584, 1268–1272 (2010).

49. Y. Sudo, H. Terashima, R. Abe-Yoshizumi, S. Kojima, M. Homma, Comparative study of the ion flux pathway in stator units of proton- and sodium-driven flagellar motors. BIOPHYSICS 5, 45–52 (2009).

50. T. Terauchi, H. Terashima, S. Kojima, M. Homma, A conserved residue, PomB-F22, in the transmembrane segment of the flagellar stator complex, has a critical role in conducting ions and generating torque. Microbiology (Reading, England) 157, 2422–2432 (2011).

51. Y. Onoue et al., Essential ion binding residues for Na+ flow in stator complex of the Vibrio flagellar motor. Scientific reports 9, 11216–11216 (2019).

52. Y. Suzuki et al., Effect of the MotA(M206I) Mutation on Torque Generation and Stator Assembly in the Salmonella H+-Driven Flagellar Motor. Journal of bacteriology 201, 19 (2019).

53. C.-J. Lo, Y. Sowa, T. Pilizota, R. M. Berry, Mechanism and kinetics of a sodium-driven bacterial flagellar motor. Proceedings of the National Academy of Sciences of the United States of America 110, E2544–2551 (2013).

54. D. F. Blair, Flagellar movement driven by proton translocation. FEBS letters 545, 86–95 (2003).

55. M. T. Mazhab-Jafari et al., Atomic model for the membrane-embedded VO motor of a eukaryotic V-ATPase. Nature 539, 118–122 (2016).

56. K. R. Baker, K. Postle, Mutations in Escherichia coli ExbB transmembrane domains identify scaffolding and signal transduction functions and exclude participation in a proton pathway. Journal of bacteriology 195, 2898–2911 (2013).

57. Y. Chang et al., Structural insights into flagellar stator-rotor interactions. eLife 8, 147 (2019).

58. D. Stock, K. Namba, L. K. Lee, Nanorotors and self-assembling macromolecular machines: the torque ring of the bacterial flagellar motor. Current Opinion in Biotechnology 23, 545–554 (2012).

59. B. L. Carroll et al., The flagellar motor of Vibrio alginolyticus undergoes major structural remodeling during rotational switching. bioRxiv 69, 2020.2004.2024.060053 (2020).

60. D. C. Fung, H. C. Berg, Powering the flagellar motor of Escherichia coli with an external voltage source. Nature 375, 809–812 (1995).

61. W. S. Ryu, R. M. Berry, H. C. Berg, Torque-generating units of the flagellar motor of Escherichia coli have a high duty ratio. Nature 403, 444–447 (2000).

62. L. D. B. Evans, C. Hughes, G. M. Fraser, Building a flagellum outside the bacterial cell. Trends in Microbiology 22, 566–572 (2014).

63. L. Pravda et al., MOLEonline: a web-based tool for analyzing channels, tunnels and pores (2018 update). Nucleic acids research 46, W368–W373 (2018).

