## Supplementary Materials for "Structure and function of stator units of the bacterial flagellar motor"

**This PDF file includes:**

Materials and Methods
Figures S1 to S21
Tables S1 to S2
References (64 – 88)

**Other Supplementary Materials for this manuscript include the following:**

Movies S1 to S5

#### Materials and Methods
##### Cloning, expression and purification

The *Cj*MotAB fragment was amplified from *Campylobacter jejuni* (ATCC BAA-2151), *So*MotAB from *Shewanella oneidensis* (ATCC 700550) and *Va*PomAB from *Vibrio alginolyticus* (ATCC 17749). They were all cloned into a modified pET vector containing a C-terminal human rhinovirus (HRV) 3C protease cleavage site and a twin-Strep-tag II (resulting in pET11a-MotA-MotB-3C-TSII). The equivalent *Se*MotAB construct complemented a Δ*motAB* mutant in *S. enterica* (fig. S2). All complexes were expressed in *E. coli* Overexpress™ C43(DE3) cells (LuBioScience GmbH) adapting published protocols (*64*). Briefly, cells were cultured in 2 l LB medium (supplemented with 100 µg/ml ampicillin) at 37 °C and protein expression was induced with 0.5 mM IPTG at $OD_{600}$ 0.4–0.8. Cells were incubated for another 3 hours at 30 °C and then harvested. The cell pellet was resuspended in 50 ml 200 mM Tris-HCl pH 8.0 and incubated at room temperature for 20 min with shaking. To gently disrupt the cells, 24.3 ml of 200 mM Tris-HCl pH 8.0 containing 1 M sucrose and 1 mM EDTA was added first, followed by addition of 330 µl 10 mg/ml lysozyme and 48 ml deionized water. After 20 min shaking at room temperature, spheroplasts were sedimented at 18,000 × *g* for 30 min. The pellet was resuspended in 100 ml 10 mM Tris-HCl pH 8.0 and incubated at room temperature for 30 min while stirring. DNase I was added to improve pellet solubilization. Membranes were then sedimented at 18,000 × *g* for 30 min, resuspended in a buffer containing 10 mM Tris-HCl pH 8.0 and 5% glycerol and stored at -80 °C. Membranes were solubilized in 1% (w:v) Lauryl Maltose Neopentyl Glycol (LMNG) (Anatrace) for 2 hours shaking on a rocking platform and then ultracentrifuged for 30 min at 100,000 × *g*. The supernatant was added to a gravity flow column containing 2 ml of Strep beads (IBA), pre-equilibrated with wash buffer (20 mM Tris-HCl pH 8.0, 150 mM NaCl, 20% glycerol and 0.005% LMNG). Beads were washed five times with 2 column volumes of wash buffer and elution was performed six times with 0.5 column volumes of elution buffer (20 mM Tris-HCl pH 8.0, 150 mM NaCl, 20% glycerol, 0.005% LMNG and 10 mM desthiobiotin).

The protein complex was then loaded onto a Superose 6, XK 16/70 gel filtration column (GE Healthcare), which was pre-equilibrated with 20 mM Tris-HCl pH 8, 150 mM NaCl and 0.002% LMNG. The peak fractions corresponding to the protein complex were concentrated to about 0.6 mg/ml using a centrifugal filter with a PES membrane (Sartorius) and used for preparation of cryo-EM sample grids.

**Sample preparation and cryo-EM data collection**

3 µl of freshly purified sample was applied onto glow-discharged (Balzers Union dual chamber CTA 010 glow discharger) grids (Quantifoil R2/1 300 mesh Cu) and plunge-frozen into liquid ethane using a Vitrobot Mark IV (FEI, Thermo Fisher Scientific). The settings were as follows: 4°C, 100% humidity, 7 s wait time, 3 s blot time, and a blot force of 0. Movies were collected using the semi-automated acquisition program EPU (FEI, Thermo Fisher Scientific) on a Titan Krios G2 microscope operated at 300 keV paired with a Falcon 3EC direct electron detector (FEI, Thermo Fisher Scientific). Images were recorded in electron counting mode, with a calibrated pixel size of 0.832 Å and defocus range of -1 to -3 µm. Number of micrographs and total exposure values for the different datasets are summarized in table S1.

**Image processing**

Image processing of the *Cj*MotAB, *Cj*MotAB(Δ41-60, D22N) and *Va*PomAB datasets was performed using RELION 3.0 (*65*) (figs. S4, S6 and S10). Micrographs were aligned and dose-weighted using MotionCor2 (*66*) and the contrast transfer function (CTF) was estimated using

CTFFIND-4.1 (*67*). The *Cj*MotAB(Δ41-60) and *So*MotAB datasets were processed in cryoSPARC Live and cryoSPARC (*68*), respectively (figs. S5 and S9). In this case, Patch motion correction and Patch CTF estimation were performed instead.

To make a 2D template for picking, a first particle picking job was done on a random subset of around 1,000 micrographs, using Laplacian-of-Gaussian picking (RELION) or blob picker (cryoSPARC Live and cryoSPARC). Selected 2D class averages were used as templates for reference-based automated particle picking from all the micrographs. Picked particles were extracted using a box size of 64 pixels (256 pixels binned 4 times). After an initial sorting and at least 4 rounds of 2D classification, selected particles were reextracted to a box size of 256 pixels and a 3D model was created *de novo*. This initial model was low-pass filtered to 60 Å and used in a 3D classification of the particles. The best class or classes were further processed in a 3D high-resolution refinement job. In cryoSPARC the map was sharpened during the same job using a dynamic mask, while in RELION mask creation and postprocessing jobs were performed to sharpen the maps by applying a B-factor corrected for the modulation transfer function of the detector. Finally, in RELION particles were further processed using per-particle CTF refinement and Bayesian polishing (including beam tilt estimation). Local resolution estimations were also obtained within RELION.

This general approach had to be modified for the *So*MotAB and *Va*PomAB datasets. Due to the preferential orientation *So*MotAB, further 3D classification jobs were performed, together with 2D classification jobs in order to keep as many side views as possible. In the case of *Va*PomAB, the initial model had to be generated with cisTEM (*69*).

Directional resolution anisotropy, which is caused by preferential orientation, was assessed using the Remote 3DFSC Processing Server (*70*) (https://3dfsc.salk.edu/). The number of picked and final particles, map-sharpening B-factor and final map resolution values for all datasets can be found in table S1.

**Atomic model building, refinement and validation**

*De novo* model building was performed manually using Coot (*71*). Models were refined using PHENIX (*72*) real space refinement. The geometry of the structures was validated using MolProbity (*73*) (table S1). Despite trying extensively, unlike for the *Cj*MotAB and *Cj*MotAB(Δ41-60, D22N) structures, it was impossible to fit a regular α-helix in the density around *Cj*MotAB(Δ41-60) MotB chain 1 D22 while respecting the chirality of this residue. The structure does fit very well to the map when introducing cis-peptides before and after this residue, which also point the carbonyl oxygen atoms to chemically plausible directions. We further validated the atomic model of *Cj*MotAB by plotting existing biological data obtained for *E. coli* MotAB from cross-linking experiments (*74*), mutational screening (*28, 75*) or tryptophan scanning analysis (*35, 36*) on the structure. Only residues above an alignment score cutoff of 2 (out of 11) for the alignment were plotted.

**Homology modeling, modeling the MotA-FliG interaction and ion channel prediction**

*Se*MotAB was modelled based on the structure of *Cj*MotAB with Modeller (*76*). The alignment (fig. S1) between *Cj*MotA and *Se*MotA, and between *Cj*MotB and *Se*MotB, respectively, was provided to model the complete *Se*MotAB heteroheptamer. Non-conserved regions (*Se*MotA 107-123 and 279-295) were removed for displaying.

*C. jejuni* FliG (*Cj*FliG residues 115-334) was modelled based on the structure of *H. pylori* $FliG_{MC1}$ (*77*) (PDB ID 3USW). The FliG $Helix_{Torque}$ peptide (*Cj*FliG residues 290-305) was extracted from this model for the following. Two subunits of MotA were extracted from the *Cj*MotAB atomic model generated in the present study and linked together from the C-terminal end of one to the N-terminal end of the other to allow submission to the FlexPepDock server (*78, 79*). The peptide was located in different positions, either on top of one MotA or in between two MotA subunits trying to maximize interaction between genetically interacting, oppositely charged residues (as described (*23*)) and uploaded to the server. The best docking results, which corresponded to FliG $Helix_{Torque}$ interacting in between MotA monomers and also correlated with the electrostatic interaction data, were later manually readjusted in terms of rotamers, distances and position of the peptide. The process of submission and selection was repeated and the result with the best score (total: 5,981.675, rmsBB: 2.320) was chosen for plotting and analysis.

For calculation of a proton and hydronium accessible channel, the *Cj*MotAB(D41-60) model was analyzed using Mole 2.5 software (*63*). The bottleneck radius was set to 1 Å. The starting points for calculation were located along the interface between MotA chain 2 and 3, from the CI helix until the MotB chain 1, according to the mutagenesis data and the conformational changed observed in this study. In order to consider the flexibility of MotA chain 1 F186, the residue was ignored for the calculation. From the resulting channels, the one with best correlation to the biological data was selected.

##### ***Salmonella enterica*** **strains and cultivation conditions**

*Salmonella enterica* serovar Typhimurium LT2 (J. Roth) (ATCC 700720) (*S. enterica*) is one of the best-studied model systems for the function of the bacterial flagellum and was therefore used for our motility experiments. The *Se*MotAB clean deletion (Δ*motAB*) and *Se*MotA/MotB amino acid point mutants were generated in *S. enterica* LT2 using the λ-RED homologous recombination system (*80*) and pET11a-*Se*MotA-*Se*MotB-3C-TSII (constructed as described for the constructs used for cryo-EM) or *S. enterica* genomic DNA as template. The *S. enterica* generalized transducing phage P22 *HT105/1 int-201* was used in transductional crosses to transfer chromosomal fragments within *S. enterica* strains (*81*). *S. enterica* strains were grown at 30 °C or 37 °C in LB supplemented with 100 µg/ml ampicillin and 1 µM IPTG if required.

**Motility assays**

Swimming motility was determined using tryptone broth (TB)-based soft agar plates containing 0.3% agar and supplemented with 100 µg/ml ampicillin and 1 µM IPTG if required. Plates were inoculated with 2 µl overnight cultures or using a pin tool (V&P Scientific) and incubated 3–4 hours at 30 °C or 37 °C. Diameters of the motility swarm were measured using ImageJ (*82*) (NIH) and normalized to the wildtype.

**Growth assays**

*S. enterica* overnight cultures were diluted 1:100 in 96-well plates and the $OD_{600}$ was measured in a microplate reader (Tecan) every 10 min for 8 hours with a brief shaking interval before each measurement. Growth rates were determined using GrowthRates 4.3 (*83*) with correlation coefficient R >0.995.

**Figure preparation**

Figures were prepared using UCSF Chimera (*84*), UCSF ChimeraX (*85*), GraphPad Prism 8 (GraphPad Software) and Illustrator (Adobe). Movies were prepared with UCSF ChimeraX, Premiere Pro (Adobe) and Keynote (Apple).

### Figures

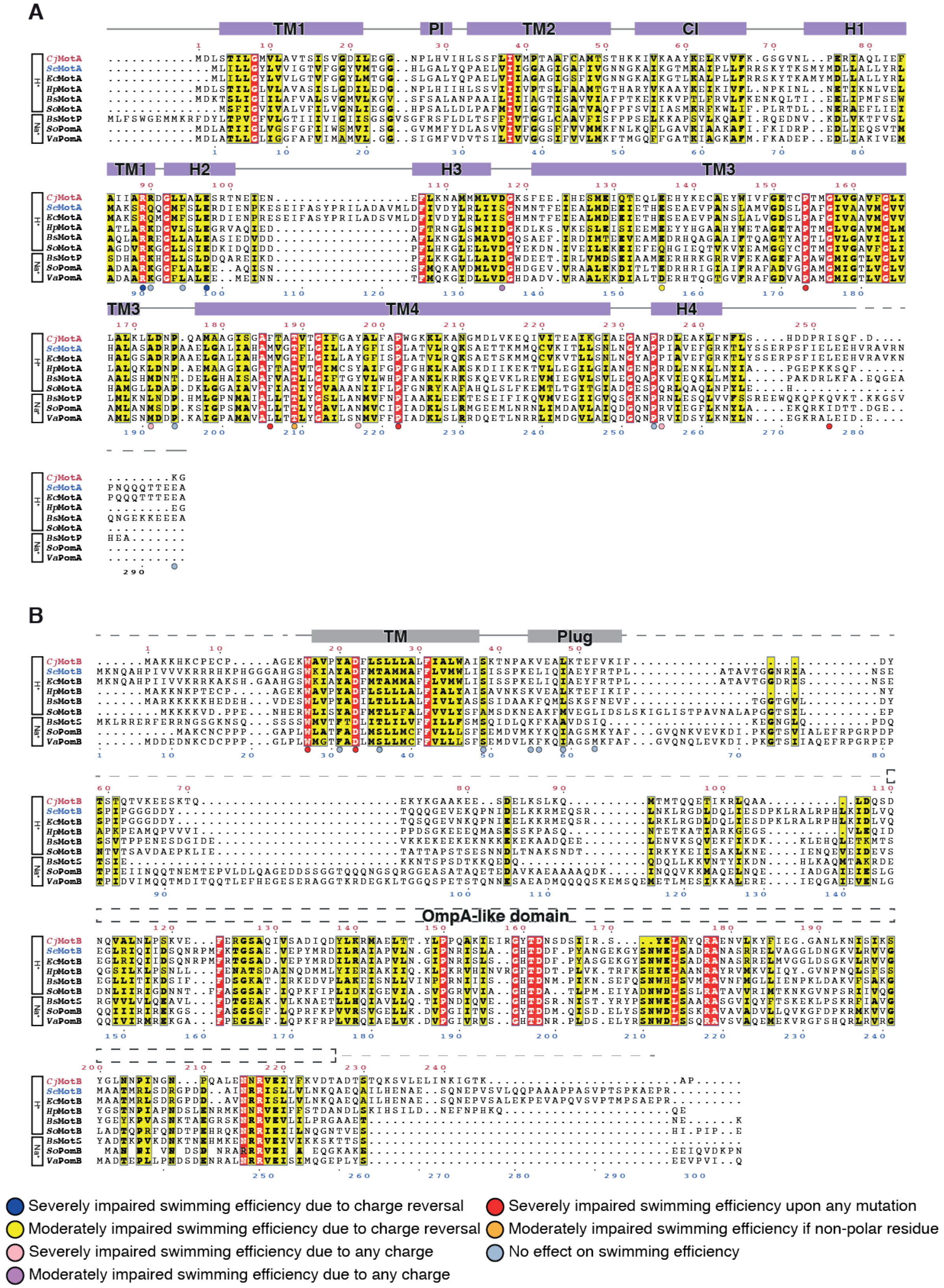
A
TM1
PI
TM2
CI
H1
H2
H3
TM3
TM4
H4
CjMotA
SeMotA
EcMotA
HpMotA
BsMotA
SoMotA
BsMotP
SoPomA
VaPomA
H+
Na+
B
TM
Plug
OmpA-like domain
CjMotB
SeMotB
EcMotB
HpMotB
BsMotB
SoMotB
BsMotS
SoPomB
VaPomB
Severely impaired swimming efficiency due to charge reversal
Moderately impaired swimming efficiency due to charge reversal
Severely impaired swimming efficiency due to any charge
Moderately impaired swimming efficiency due to any charge
Severely impaired swimming efficiency upon any mutation
Moderately impaired swimming efficiency if non-polar residue
No effect on swimming efficiency

**Fig. S1. Sequence alignment of MotA and MotB homologues in different species.** (**A** and **B**) Multiple sequence alignment of MotA (A) and MotB (B). The proteins are subdivided into proton and sodium channels. Residue numbers above the sequences (red) correspond to the *C. jejuni* residue numbers, while residue numbers below the sequences (blue) correspond to those of *S. enterica*. Residues marked with a circle indicate residues mutated in *S. enterica*. α-helices are indicated by solid boxes, the dashed lines indicate that the structure was not resolved in this study. The OmpA-like domain of MotB is also indicated above the alignment. Amino acids that are identical or partially conserved are colored red and yellow, respectively. Species abbreviations: *Cj*, *Campylobacter jejuni*; *Se*, *Salmonella enterica*; *Ec*, *Escherichia coli*; *Hp*, *Helicobacter pylori*; *Bs*, *Bacillus subtilis*; *So*, *Shewanella oneidensis*; *Va*, *Vibrio alginolyticus*.

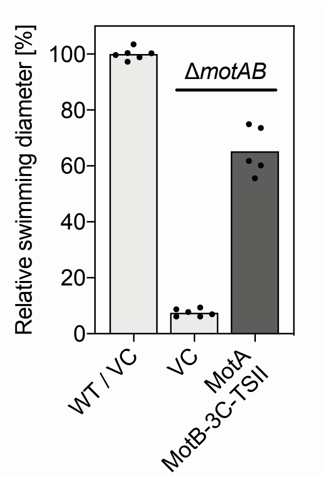

**Fig. S2. Swimming motility of C-terminal tagged *Se*MotB.** Swimming motility phenotype of *S. enterica* Δ*motAB* complemented with a plasmid expressing C-terminal tagged *Se*MotB (pET11a-MotA-MotB-3C-TwinStrepII) using soft agar motility plates containing 0.3% agar. Expression of *Se*MotA and *Se*MotB with C-terminal twin-Strep-tag II from pET11a-MotA-MotB-3C-TwinStrepII was induced by addition of 1 µM IPTG. The diameters of the motility swarm were measured and normalized to the wildtype containing the empty vector. The bar graphs represent the mean of at least five biological replicates. WT, wildtype; VC, empty vector control; 3C-TSII, C-terminal HRV 3C protease-cleavable twin-Strep-tag II.

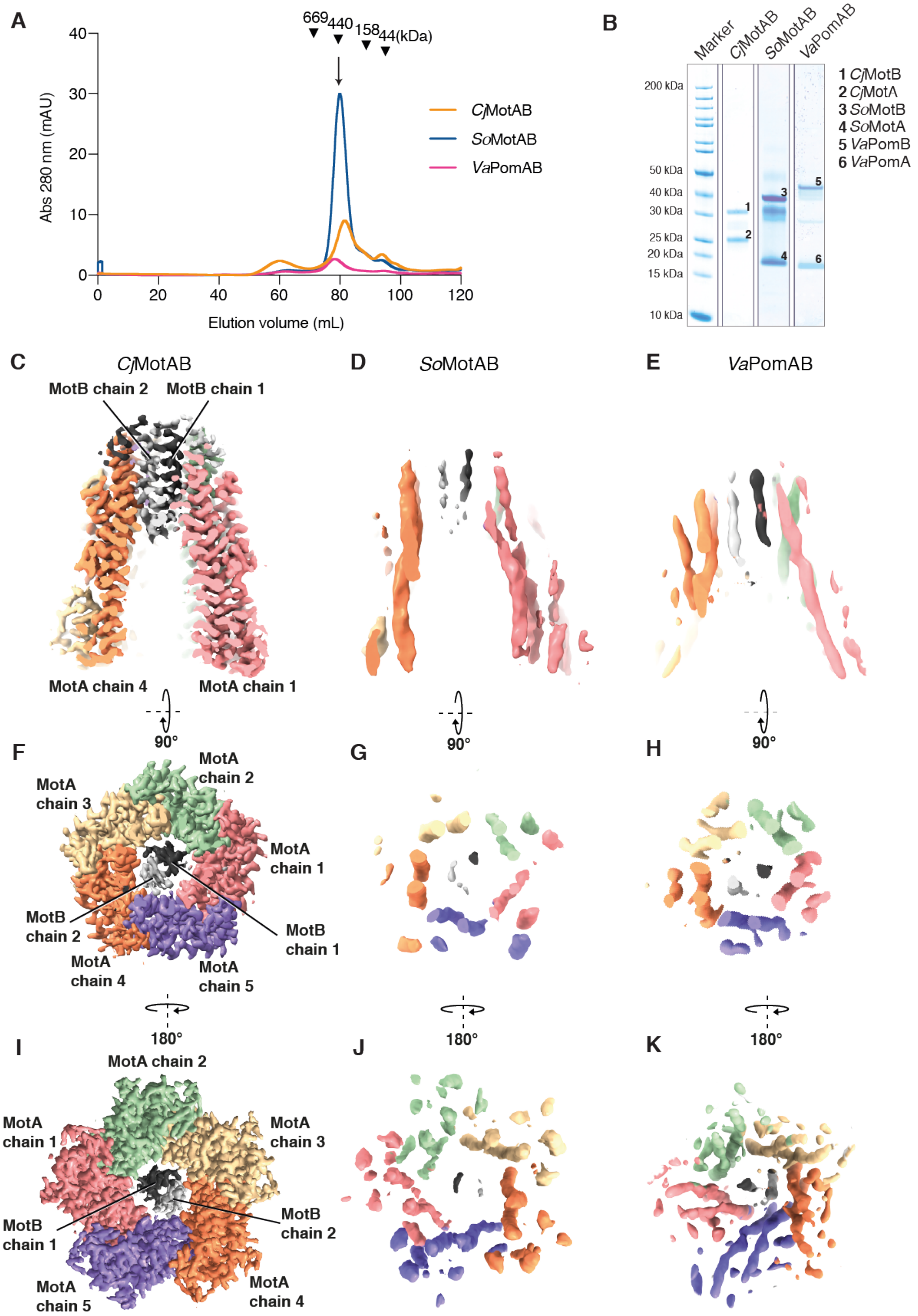
A
669
440
158
44(kDa)
Abs 280 nm (mAU)
Elution volume (mL)
CjMotAB
SoMotAB
VaPomAB
B
Marker
CjMotAB
SoMotAB
VaPomAB
200 kDa
50 kDa
40 kDa
30 kDa
25 kDa
20 kDa
15 kDa
10 kDa
1 CjMotB
2 CjMotA
3 SoMotB
4 SoMotA
5 VaPomB
6 VaPomA
C
CjMotAB
MotB chain 2
MotB chain 1
MotA chain 4
MotA chain 1
D
SoMotAB
E
VaPomAB
90°
F
MotA chain 2
MotA chain 3
MotA chain 1
MotB chain 2
MotB chain 1
MotA chain 4
MotA chain 5
G
H
180°
I
MotA chain 2
MotA chain 1
MotA chain 3
MotB chain 2
MotB chain 1
MotA chain 5
MotA chain 4
J
K

**Fig. S3. Purification and stoichiometry of MotAB homologues.** (**A**) Size-exclusion chromatography (SEC) profile of detergent-purified *C. jejuni* MotAB (*Cj*MotAB), *S. oneidensis* MotAB (*So*MotAB) and *V. alginolyticus* PomAB (*Va*PomAB) complexes. The fraction used for cryo-EM grid preparation is indicated with an arrow. Elution volumes of molecular weight standards are indicated with inverted triangles. (**B**) The corresponding SDS-PAGE gels for (A) are shown. (**C**, **F** and **I**) Side view, top view and bottom view of *Cj*MotAB. Same color code as in Fig. 1. (**D**, **G** and **J**) Side view, top view and bottom view of *So*MotAB. Same color code as for *Cj*MotAB. **(E**, **H** and **K**) Side view, top view and bottom view of *Va*PomAB. Same color code as for *Cj*MotAB.

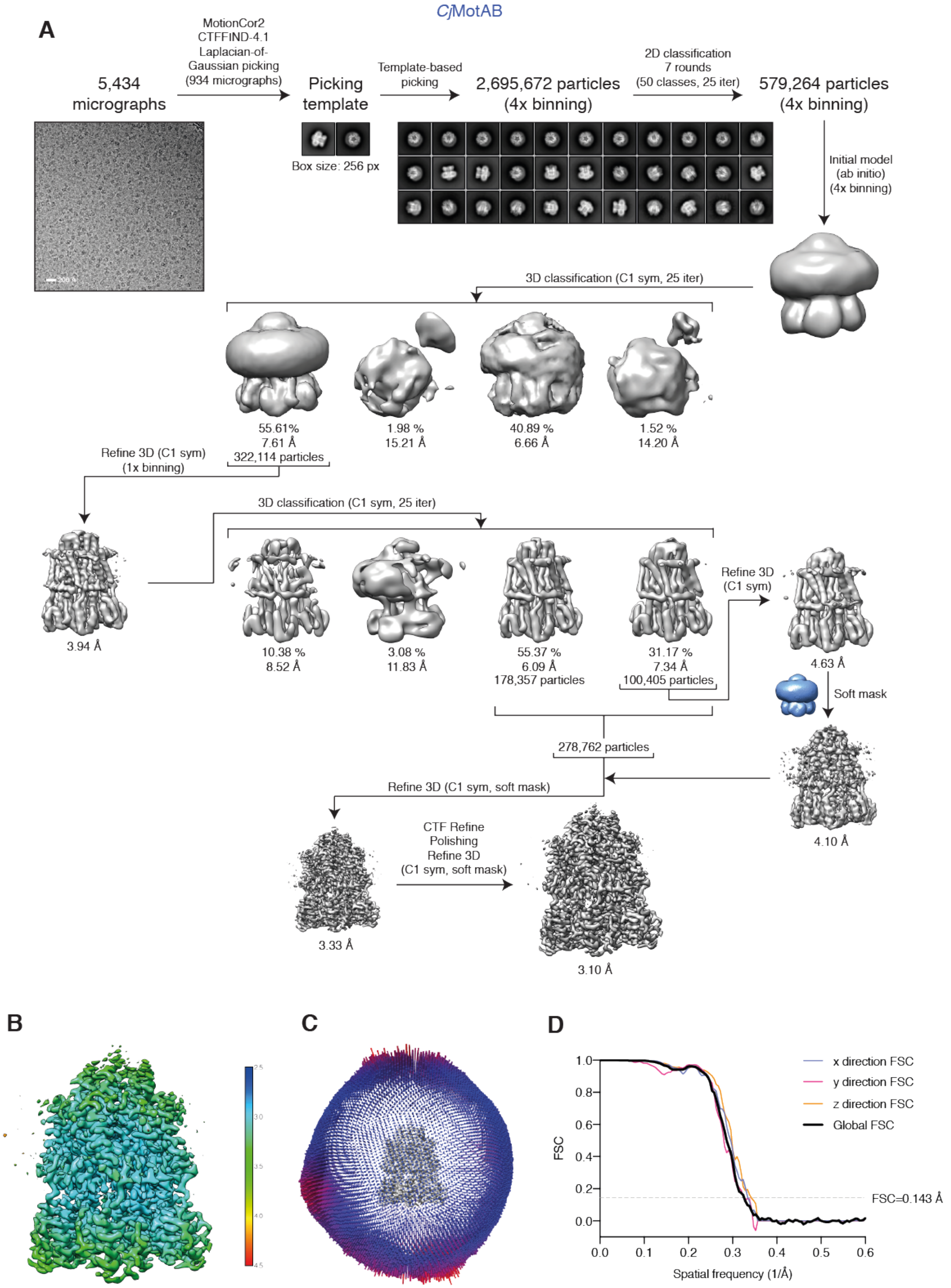

**Fig. S4. Cryo-EM of *Cj*MotAB.** (**A**) Flowchart of the data collection and processing pipeline in RELION that resulted in the final *Cj*MotAB Cryo-EM structure. 2,695,672 particles were picked from 5,434 micrographs. After 7 rounds of 2D classification, 579,264 particles were used to generate an initial model, followed by a 3D classification job with 4 classes. The best class containing 322,114 particles was unbinned and further 3D refined, obtaining a 3.94 Å resolution map. In order to improve resolution of MotB, another 3D classification was performed and class 4 was 3D refined, obtaining a 4.10 Å resolution map. To improve the resolution, particles from class 3 were also selected for another 3D refinement job. After masking, per-particle CTF refinement and Bayesian polishing, the map reached a resolution of 3.10 Å. (**B**) Cryo-EM density map of *Cj*MotAB colored by local resolution (in Å) estimated in RELION. (**C**) Euler angular distribution plotting for *Cj*MotAB. (**D**) 3D Fourier shell correlation (FSC) curves for *Cj*MotAB. Global resolution is estimated to be 3.10 Å at FSC=0.143 (dashed line).

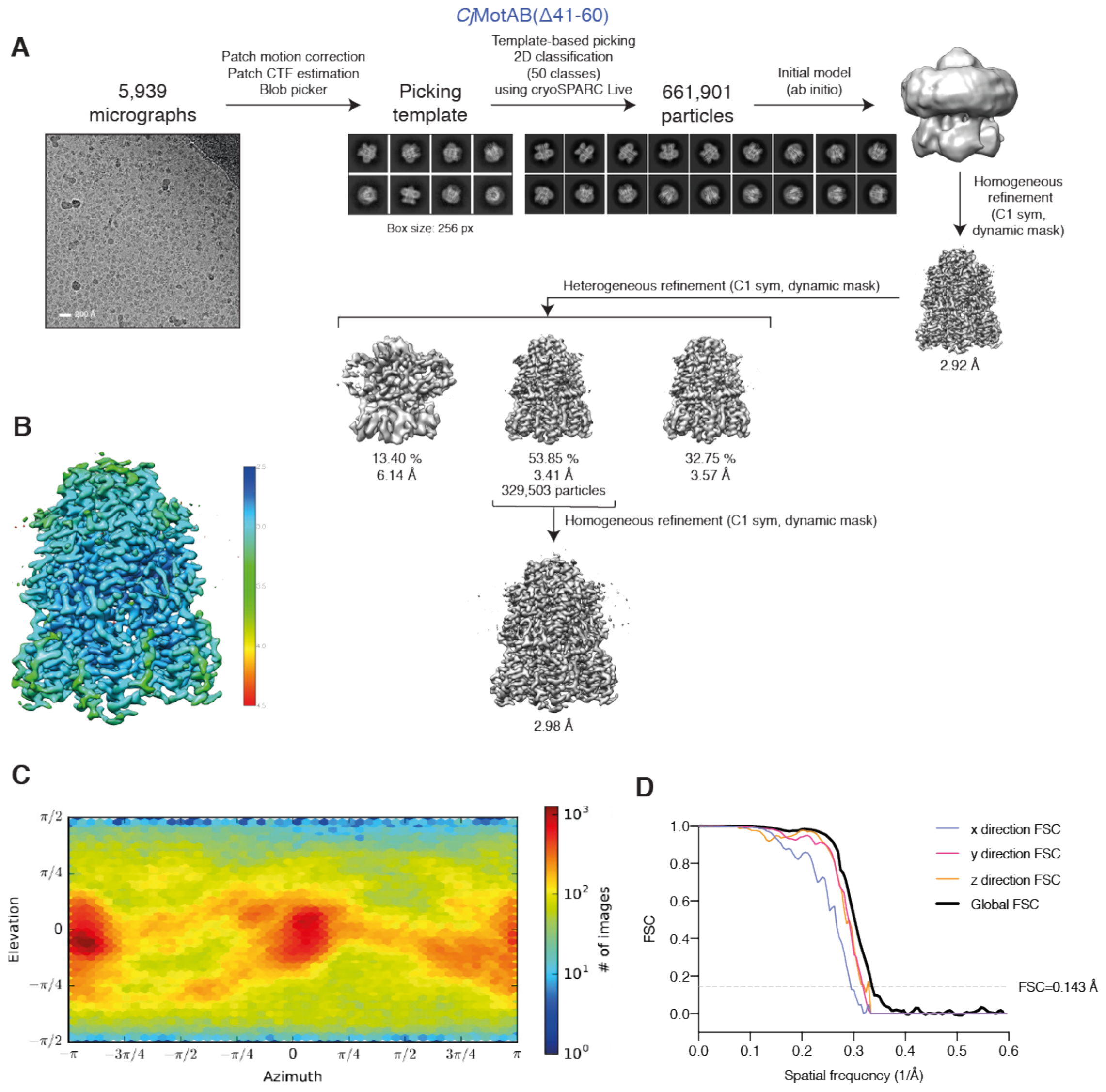

**Fig. S5. Cryo-EM of *Cj*MotAB(Δ41-60).** (**A**) Flowchart of the data collection and processing pipeline in cryoSPARC live that resulted in the final *Cj*MotAB(Δ41-60) cryo-EM structure. 5,939 micrographs were processed on-the-fly, and the 661,901 particles selected gave a 2.92 Å resolution map. In order to improve resolution of the TM domain of MotB, a heterogeneous refinement job was performed, and the best resulting volume (329,503 particles) was further refined, achieving a resolution of 2.98 Å. (**B**) Cryo-EM density map of *Cj*MotAB(Δ41-60) colored by local resolution (in Å) estimated in cryoSPARC. (**C**) Euler angular distribution plotting for *Cj*MotAB(Δ41-60)

generated in cryoSPARC. (**D**) 3D Fourier shell correlation (FSC) curves for *Cj*MotAB(Δ41-60). Global resolution is estimated to be 2.98 Å at FSC=0.143 (dashed line).

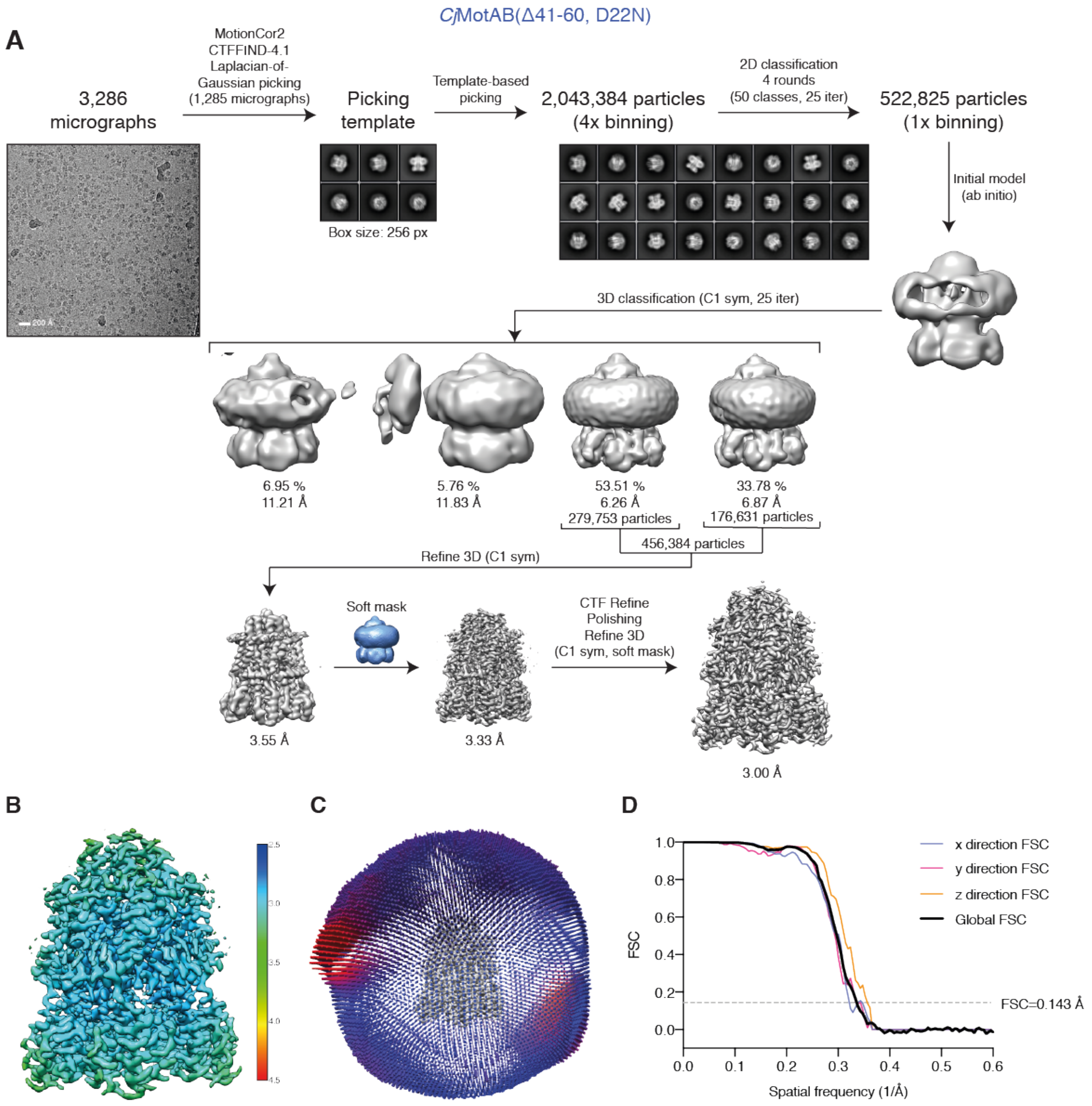

**Fig. S6. Cryo-EM of *Cj*MotAB(Δ41-60, D22N).** (**A**) Flowchart of the data collection and processing pipeline in RELION that resulted in the final *Cj*MotAB(Δ41-60, D22N) cryo-EM structure. 2,043,384 particles were picked from 3,286 micrographs. After 4 rounds of 2D classification, 522,825 particles were unbinned and used to generate an initial model and a following 3D classification with 4 classes. The best 2 classes (456,384 particles) were further 3D refined, obtaining a 3.55 Å resolution map. After masking, per-particle CTF refinement and Bayesian polishing, the map reached a resolution of 3.00 Å. (**B**) Cryo-EM density map of

*Cj*MotAB(Δ41-60, D22N) colored by local resolution (in Å) estimated in RELION. (**C**) Euler angular distribution plotting for *Cj*MotAB(Δ41-60, D22N). (**D**) 3D Fourier shell correlation (FSC) curves for *Cj*MotAB(Δ41-60, D22N). Global resolution is estimated to be 3.00 Å at FSC=0.143 (dashed line).

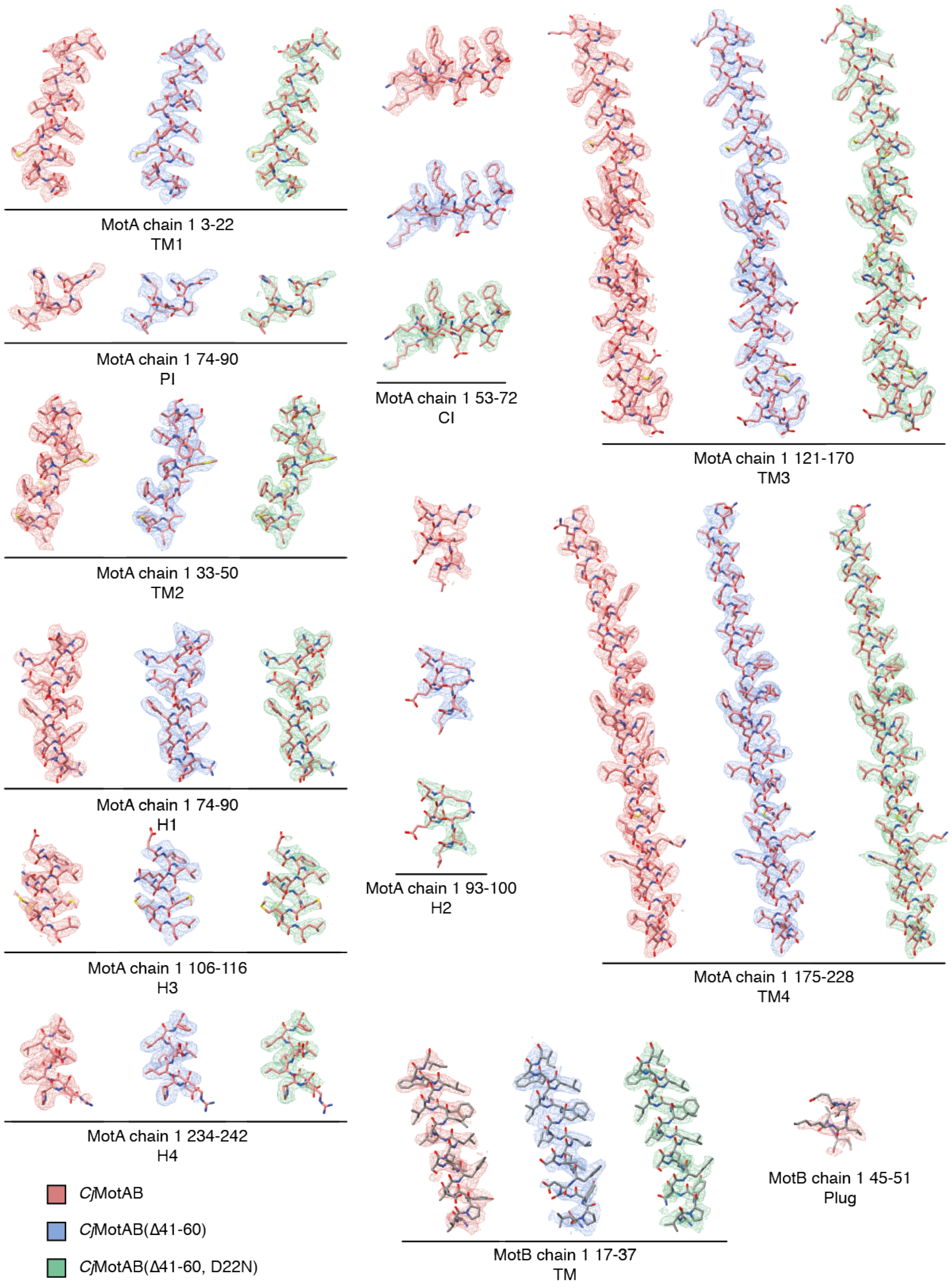

**Fig. S7. Fit of the *Cj*MotAB atomic models to the cryo-EM maps.** Fragments of atomic models and the corresponding fragments of electrostatic potential maps, of the α-helices of *Cj*MotA and

*Cj*MotB for the *Cj*MotAB (red), *Cj*MotAB(Δ41-60) (blue) and *Cj*MotAB(Δ41-60, D22N) (green) structures.

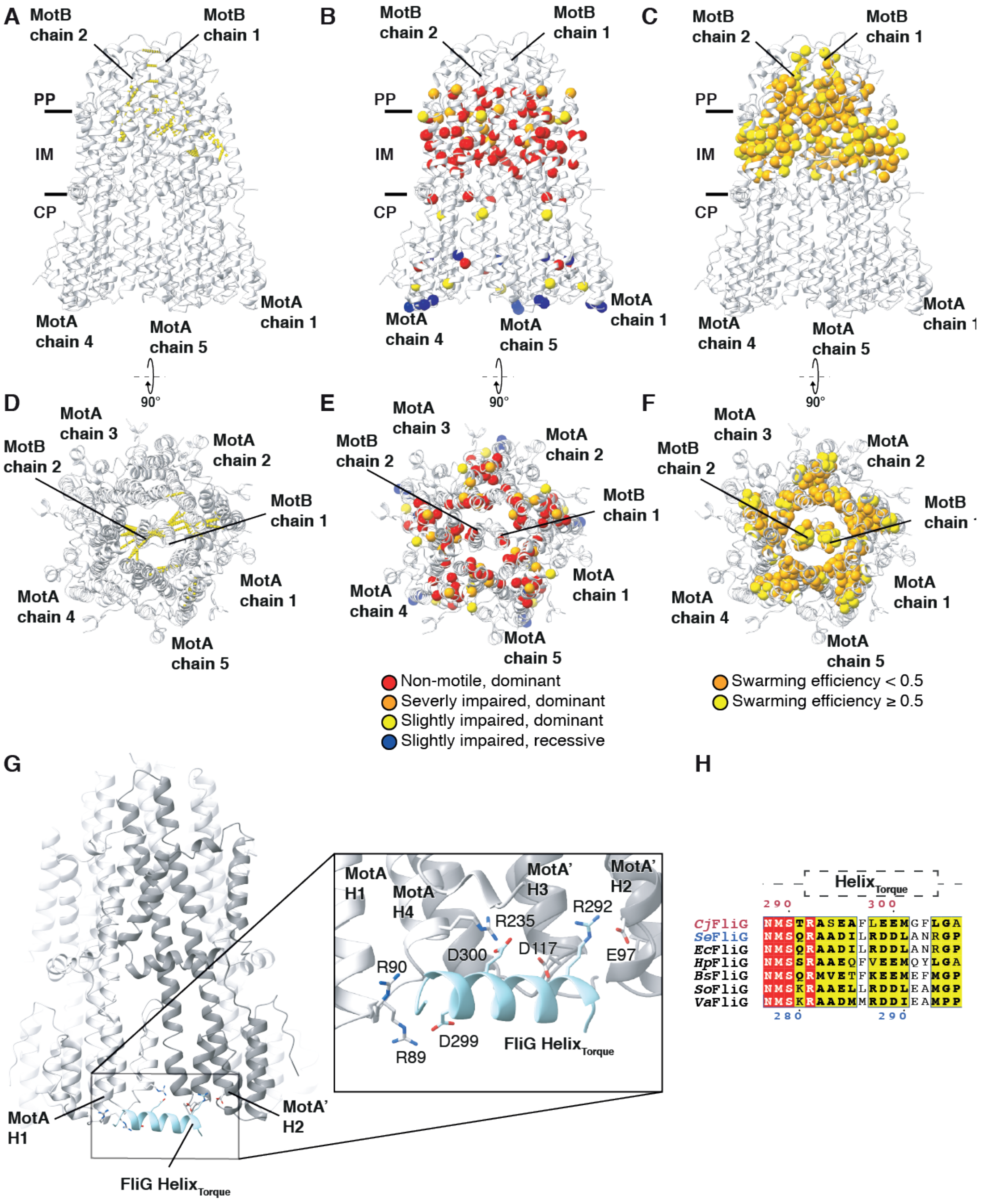

**Fig. S8. Validation of *Cj*MotAB structure by prior functional data and modelling of MotA-FliG interaction.** (A to C) Side views and (D to F) top views of the validation of the *Cj*MotAB structure, see Methods for details. (**A** and **D**) Representation (shown as dotted yellow lines between the homologous $C_\alpha$ atoms) of *E. coli* MotAB (*Ec*MotAB) residues that can be crosslinked when they are mutated to cysteines (*74*), mapped on the *Cj*MotAB structure. Of all possible crosslinks, only the result closest to 5 Å is displayed. A slice of the protein structure is shown. Note that the observed crosslinks are consistent with the expected $C_\alpha$ distances of cysteine crosslinks (5 Å), and that the crosslinks confirm the register of the MotB helices. (**B** and **E**) Mutational analysis for *Ec*MotA (*28*) and *Ec*MotB (*75*) plotted onto the *Cj*MotAB structure (represented as spheres on the position of the homologous $C_\alpha$ atom), colored by severity of the phenotype. Observe the distribution of phenotype severity according to the position in the structure. (**C** and **F)** Results of tryptophan scanning analyses for *Ec*MotA (*35*) and *Ec*MotB (*36*), plotted onto the *Cj*MotAB structure, represented as spheres on the position of the homologous $C_\alpha$ atom and colored by their impact on the swarming efficiency of *E. coli*. Note the different distribution of residues that can be more easily mutated to tryptophan (swarming efficiency > 0.5) (e.g. because they are interacting with the aliphatic chains of membrane lipids), vs. those that cannot (swarming efficiency <0.5) (e.g. because they are buried in the structure). (**G**) Modeling of the interaction between *Cj*MotA and the *Cj*FliG torque helix ($Helix_{Torque}$) (see Methods) showing the interaction between the residues D299 and R292 of FliG with the residues R89 of one MotA subunit (light grey) and the residue E97 of the adjacent MotA subunit (light grey; MotA'). $Helix_{Torque}$: light blue. Charged residues on the interaction surface are shown in stick representation. (**H**) Conservation of FliG $Helix_{Torque}$. Top position numbers (red) refer to the *C. jejuni* FliG sequence, while bottom position numbers (blue) correlate with *S. enterica* FliG. $Helix_{Torque}$ is indicated by a dotted-line box. Amino acids that are identical or partially conserved are colored red and yellow, respectively. Species abbreviations: *Cj*, *Campylobacter jejuni*; *Se*, *Salmonella enterica*; *Ec*, *Escherichia coli*; *Hp*, *Helicobacter pylori*; *Bs*, *Bacillus subtilis*; *So*, *Shewanella oneidensis*; *Va*, *Vibrio alginolyticus*.

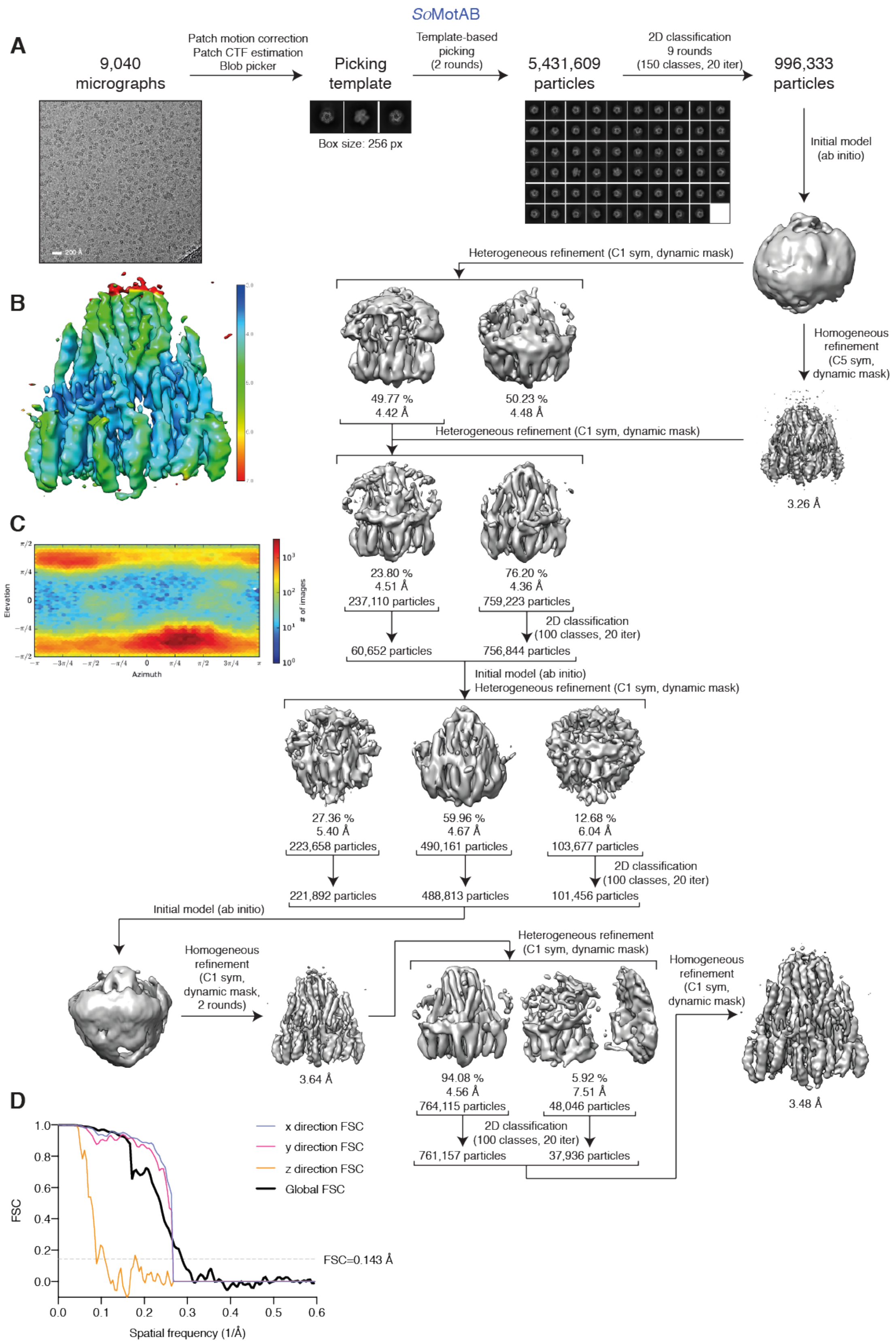

SoMotAB
A
9,040 micrographs
Patch motion correction
Patch CTF estimation
Blob picker
Picking template
Box size: 256 px
Template-based picking (2 rounds)
5,431,609 particles
2D classification 9 rounds (150 classes, 20 iter)
996,333 particles
Initial model (ab initio)
200 Å
Heterogeneous refinement (C1 sym, dynamic mask)
B
Homogeneous refinement (C5 sym, dynamic mask)
49.77 %
4.42 Å
50.23 %
4.48 Å
Heterogeneous refinement (C1 sym, dynamic mask)
3.26 Å
C
23.80 %
4.51 Å
237,110 particles
76.20 %
4.36 Å
759,223 particles
2D classification (100 classes, 20 iter)
60,652 particles
756,844 particles
Elevation
Azimuth
### of images
Initial model (ab initio)
Heterogeneous refinement (C1 sym, dynamic mask)
27.36 %
5.40 Å
223,658 particles
59.96 %
4.67 Å
490,161 particles
12.68 %
6.04 Å
103,677 particles
2D classification (100 classes, 20 iter)
221,892 particles
488,813 particles
101,456 particles
Initial model (ab initio)
Heterogeneous refinement (C1 sym, dynamic mask)
Homogeneous refinement (C1 sym, dynamic mask, 2 rounds)
Homogeneous refinement (C1 sym, dynamic mask)
3.64 Å
94.08 %
4.56 Å
764,115 particles
5.92 %
7.51 Å
48,046 particles
3.48 Å
D
2D classification (100 classes, 20 iter)
761,157 particles
37,936 particles
x direction FSC
y direction FSC
z direction FSC
Global FSC
FSC
FSC=0.143 Å
Spatial frequency (1/Å)

**Fig. S9. Cryo-EM *So*MotAB.** (**A**) Flowchart of the data collection and processing pipeline in cryoSPARC that resulted in the final *So*MotAB Cryo-EM structure. 5,431,609 particles were picked from 9,040 micrographs. After 9 rounds of 2D classification, 996,333 particles were used to generate an initial model, which was used for both homogenous and heterogeneous refinement. The resulting map from the homogenous refinement and the first class from the heterogeneous refinement were used as initial volumes for another heterogenous refinement with all particles. After 2D classification was applied to both classes, particles from the best 2D classes were pooled together for another round heterogenous refinement. 2D classification was performed again to each class, and best 2D classes pooled together to obtain a new initial model. Another homogenous refinement job was performed, followed by a heterogenous refinement and 2D classification of each class, in order to keep all the 2D classes with side views. The final map with 799,093 particles reached a resolution of 3.48 Å. (**B**) Cryo-EM density map of *So*MotAB colored by local resolution (in Å) estimated in cryoSPARC. **C**, Euler angular distribution plotting for *So*MotAB generated in cryoSPARC. (**D**) 3D Fourier shell correlation (FSC) curves for *So*MotAB. Global resolution is estimated to be 3.48 Å (cryoSPARC) at FSC=0.143 (dashed line).

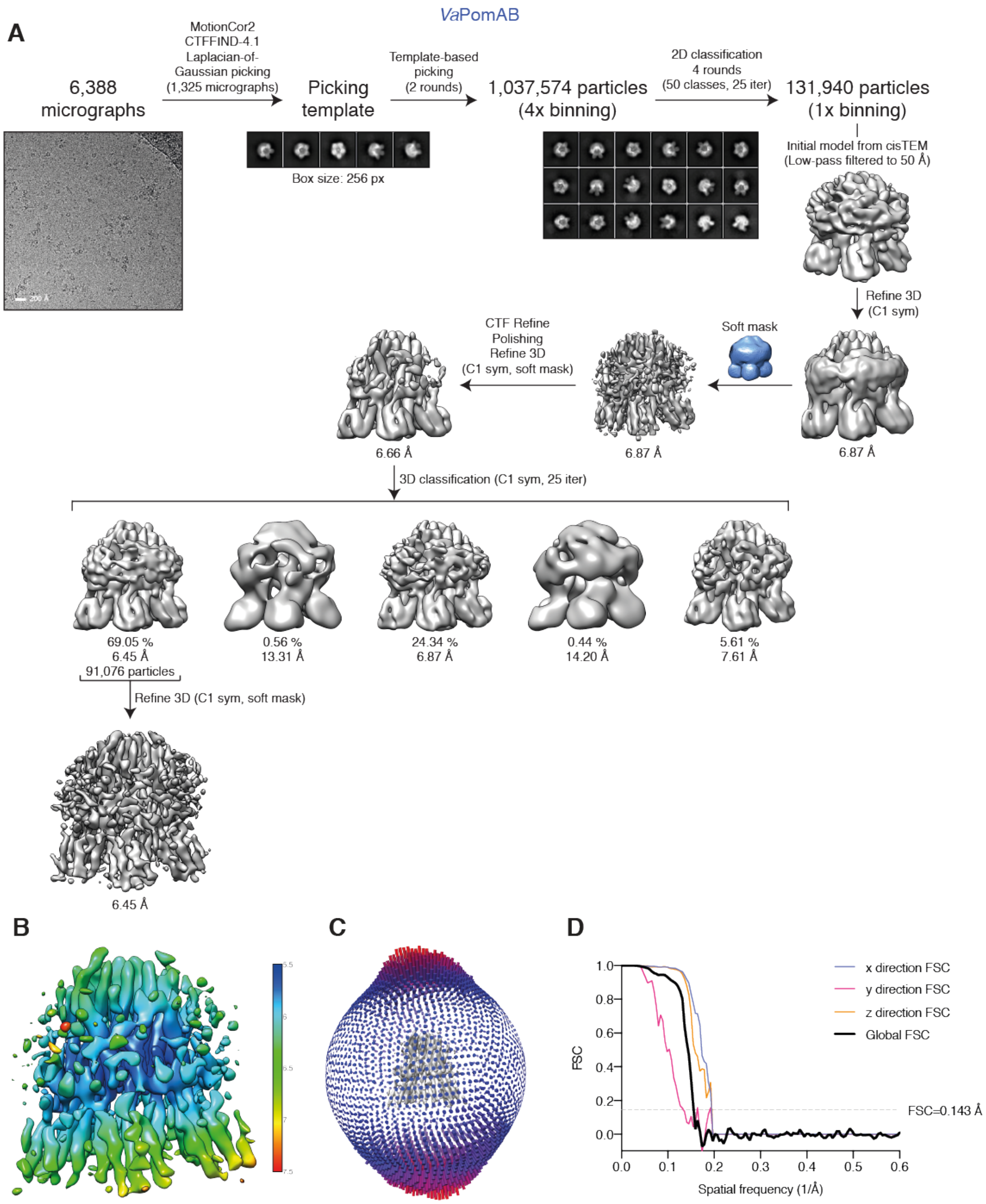

**Fig. S10. Cryo-EM *Va*PomAB.** (**A**) Flowchart of the data collection and processing pipeline in RELION that resulted in the final *Va*PomAB Cryo-EM structure. 1,037,574 particles were picked from 6,388 micrographs. After 4 rounds of 2D classification, 131,940 particles were selected and further processed in a 3D refinement job, using an initial model generated in cisTEM. This resulted in a 6.66 Å resolution volume (after masking, per-particle CTF refinement and Bayesian polishing). To resolve heterogeneity and to improve resolution, a 3D classification was performed. The first class (91,076 particles) was further refined obtaining a 6.45 Å resolution map. (**B**) Cryo-EM density map of *Va*PomAB colored by local resolution (in Å) estimated in RELION. (**C**) Euler angular distribution plotting for *Va*PomAB. (**D**) 3D Fourier shell correlation (FSC) curves for *Va*PomAB. Global resolution is estimated to be 6.45 Å at FSC=0.143 (dashed line).

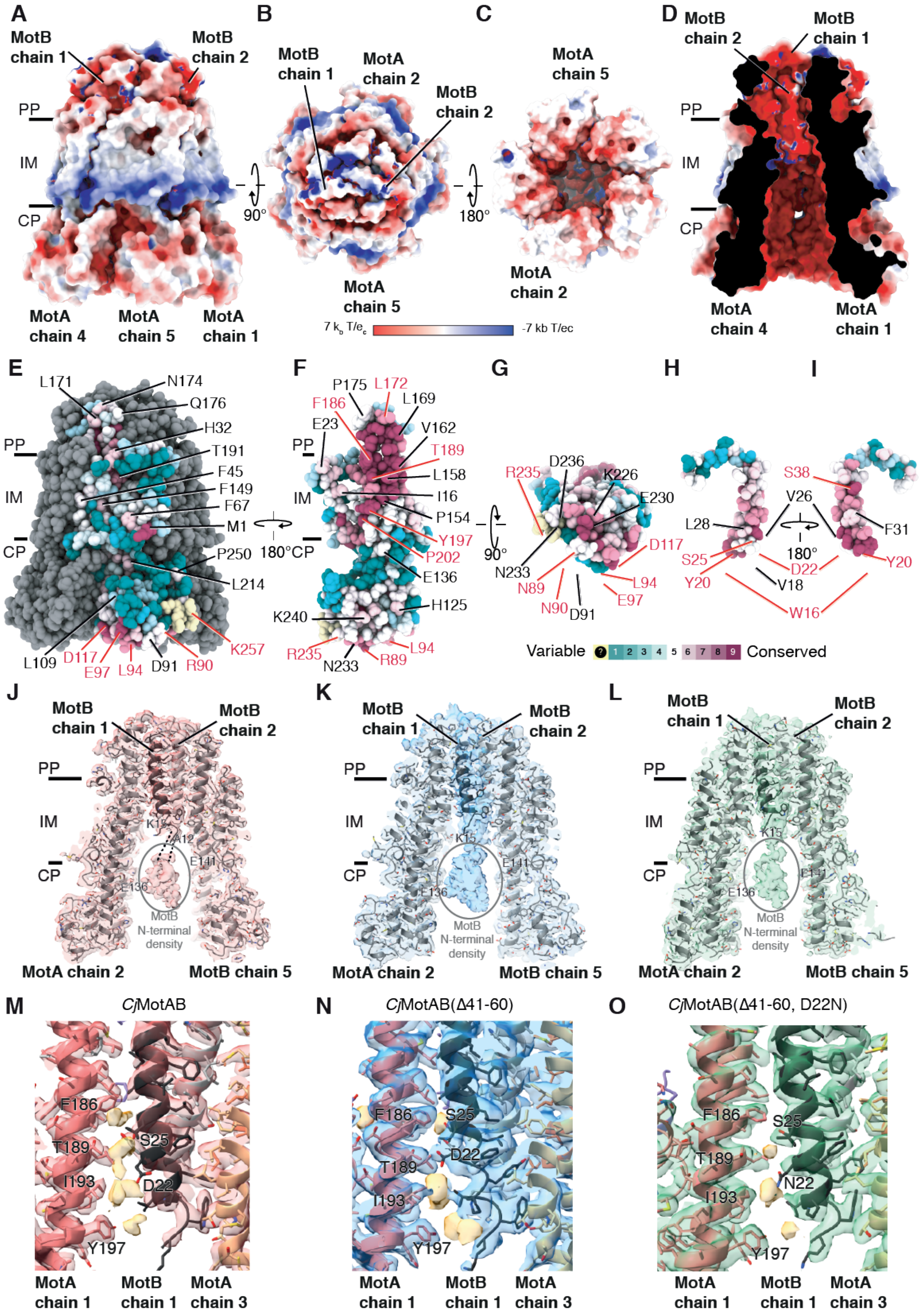

A
MotB chain 1
MotB chain 2
PP
IM
CP
MotA chain 4
MotA chain 5
MotA chain 1
B
MotB chain 1
MotA chain 2
MotB chain 2
90°
MotA chain 5
C
MotA chain 5
180°
MotA chain 2
7 kb T/ec
-7 kb T/ec
D
MotB chain 2
MotB chain 1
PP
IM
CP
MotA chain 4
MotA chain 1
E
L171
N174
Q176
H32
T191
F45
F149
F67
M1
P250
L214
PP
IM
CP
L109
D117
E97
L94
D91
R90
K257
180°
F
P175
L172
L169
F186
E23
V162
T189
L158
I16
P154
Y197
P202
E136
H125
K240
R235
N233
R89
L94
PP
IM
CP
G
90°
R235
D236
K226
E230
D117
N233
N89
N90
D91
L94
E97
H
I
S38
V26
L28
F31
180°
S25
D22
Y20
V18
Y20
W16
Variable
Conserved
J
MotB chain 1
MotB chain 2
PP
IM
CP
K15
A12
E141
E136
MotB N-terminal density
MotA chain 2
MotB chain 5
K
MotB chain 1
MotB chain 2
PP
IM
CP
K15
E141
E136
MotB N-terminal density
MotA chain 2
MotB chain 5
L
MotB chain 1
MotB chain 2
PP
IM
CP
K15
E141
E136
MotB N-terminal density
MotA chain 2
MotB chain 5
M
CjMotAB
F186
T189
S25
I193
D22
Y197
MotA chain 1
MotB chain 1
MotA chain 3
N
CjMotAB(Δ41-60)
F186
S25
D22
T189
I193
Y197
MotA chain 1
MotB chain 1
MotA chain 3
O
CjMotAB(Δ41-60, D22N)
F186
S25
T189
N22
I193
Y197
MotA chain 1
MotB chain 1
MotA chain 3

**Fig. S11. Charge and surface conservation of *Cj*MotAB, density for the N-terminal region of *Cj*MotB and putative solvent molecules in the channel.** (**A** to **D**) Exposed surface of MotAB from the side (A), top (B), bottom (C) and as a sliced side view (D) colored according to its electrostatic surface potential, calculated with APBS (*86*). (**E** to **I**) Conservation (calculated with ConSurf (*87*)) of the surface residues of MotA from external side (E), internal side (F) and bottom (G); and of MotB from both sides (H-I). Atom representation of the model colored by conservation. All residues with a ConSurf conservation of ≥5 have been labeled. Homologous residues mutated in this study in *S. enterica* are labeled in red. (**J** to **L**) Representation of the *Cj*MotAB (J), *Cj*MotAB(Δ41-60) (K), and *Cj*MotAB(Δ41-60, D22N) (L) electrostatic potential maps at low threshold, together with the ribbon model representation with side chains of the corresponding atomic models, to illustrate the non-modeled density corresponding to the MotB N-terminal region. (**M** to **O**) Representation of *Cj*MotAB (M), *Cj*MotAB(Δ41-60) (N), and *Cj*MotAB(Δ41-60, D22N) (O) channels and putative solvent molecules. Density not modeled with atomic model, which is thought to correspond to solvent molecules (such as water) has been colored in pale yellow. PP, periplasm; IM, inner membrane; CP, cytoplasm.

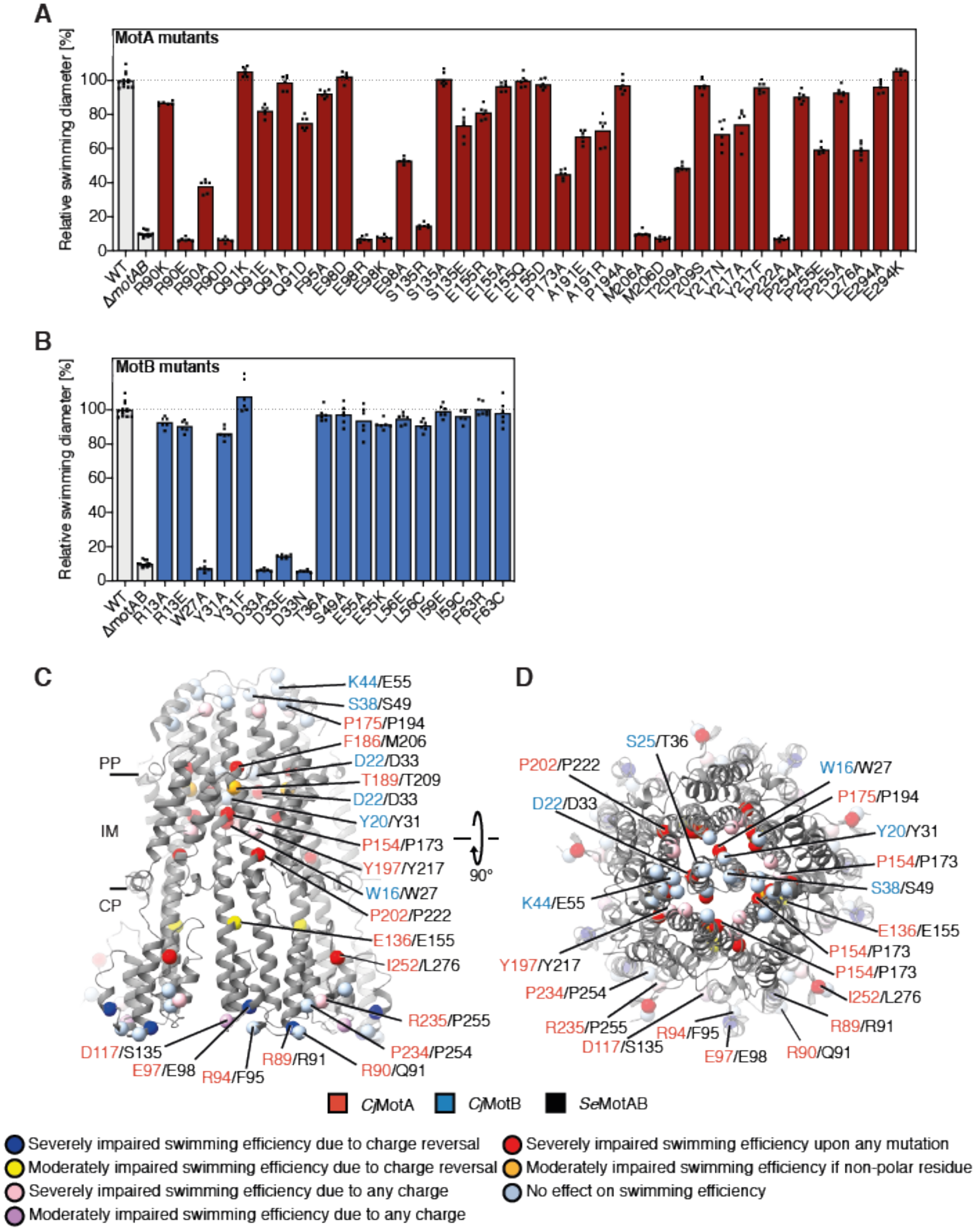

**Fig. S12. Swimming motility phenotype of *Se*MotAB point mutants. (A and B**) The motility phenotypes of *S. enterica* MotA (A) and MotB (B) point mutants were analyzed in soft-agar plates containing 0.3% agar at 37 °C. The diameters of the motility swarm were measured and normalized to the wildtype. The bar graphs represent the mean of at least six biologically independent samples. Replicates are shown as individual data points. WT, wildtype. (**C** and **D**) Swimming efficiency of the *S. enterica* point mutants, plotting the mutated residues as spheres on the *Cj*MotAB structure (grey) on the position of the $C_\alpha$ atoms of homologous residues in *C. jejuni* as side (C) and top (D) view. The corresponding *C. jejuni* residue is listed first and colored red (*Cj*MotA) or blue (*Cj*MotB), the residue number of *Se*MotA or *Se*MotB that was mutated is shown in black.

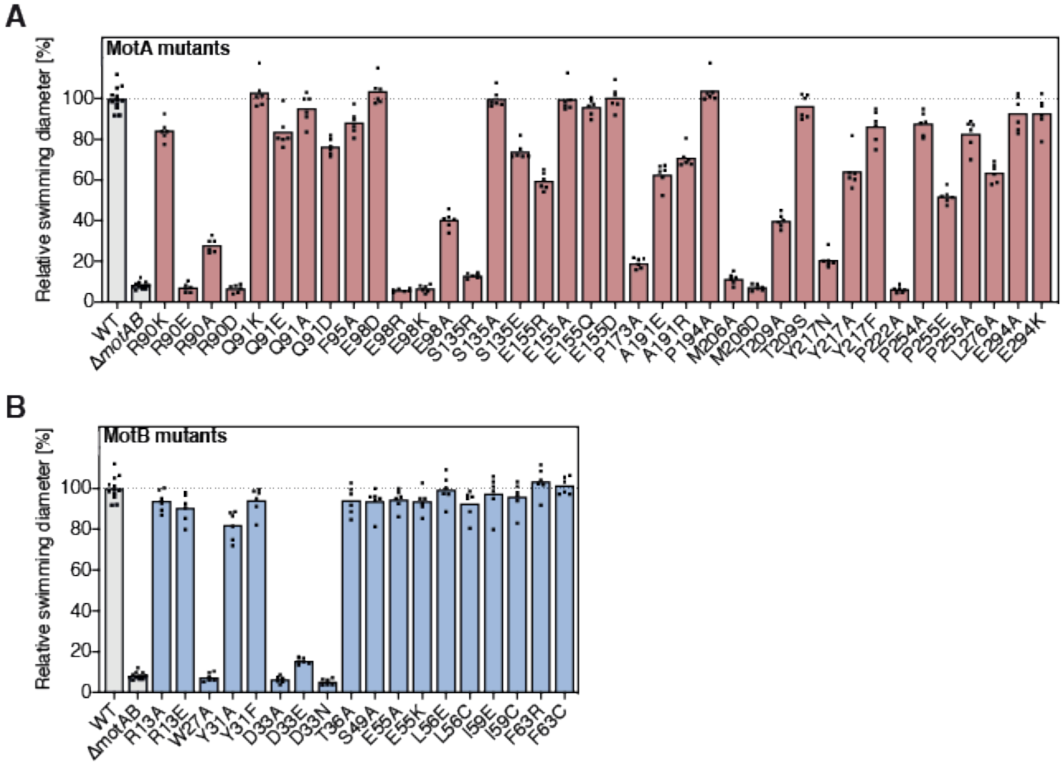

**Fig. S13. Swimming motility of *Se*MotAB point mutants at 30 °C.** (**A** and **B**) The motility phenotypes of *S. enterica* MotA (A) and MotB (B) point mutants were analyzed in soft-agar plates containing 0.3% agar. The diameters of the motility swarm were measured and normalized to the wildtype. The bar graphs represent the mean of at least six biologically independent samples. Replicates are shown as individual data points. WT, wildtype.

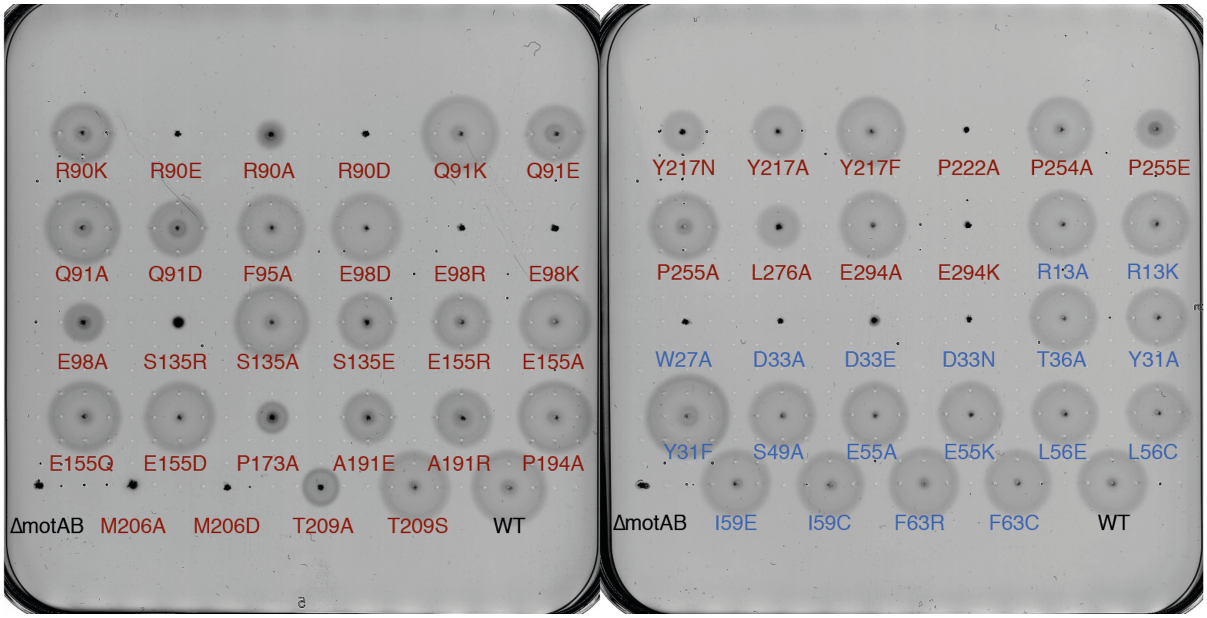

**Fig. S14. Representative swimming motility plates of *Se*MotAB mutants at 37 °C.** Exemplary soft agar plates inoculated with *S. enterica* MotA (red) and MotB (blue) point mutants in comparison to the wildtype and Δ*motAB*. WT, wildtype.

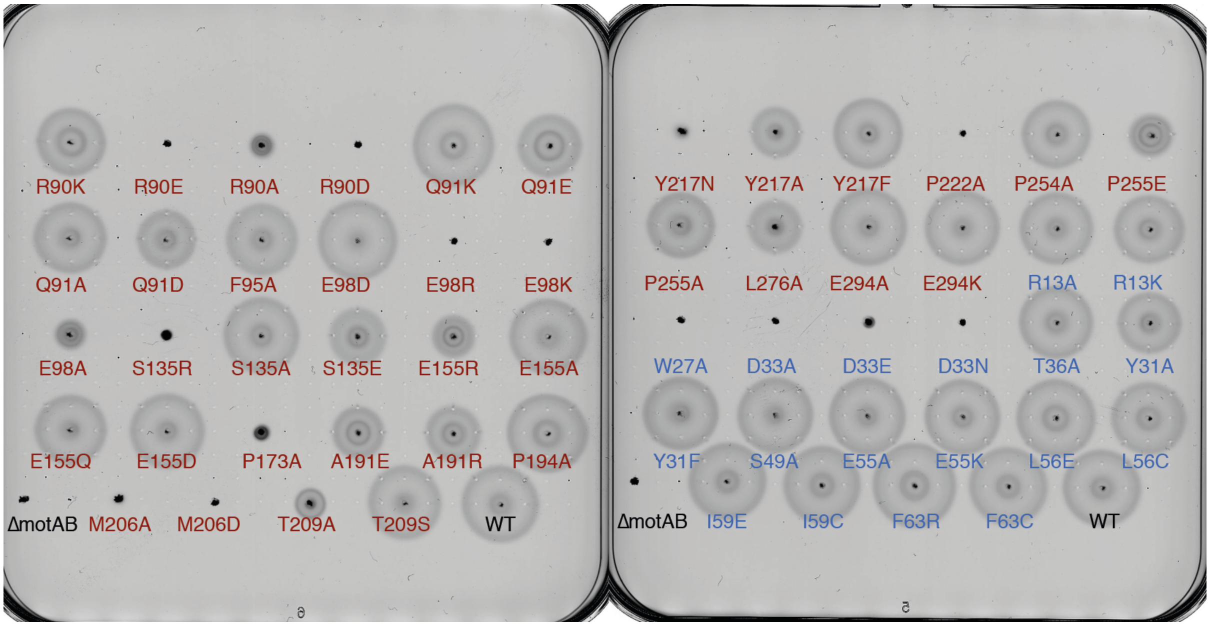

**Fig. S15. Representative swimming motility plates of *Se*MotAB mutants at 30 °C.** Exemplary soft agar plates inoculated with *S. enterica* MotA (red) and MotB (blue) point mutants in comparison to the wildtype and Δ*motAB*. WT, wildtype.

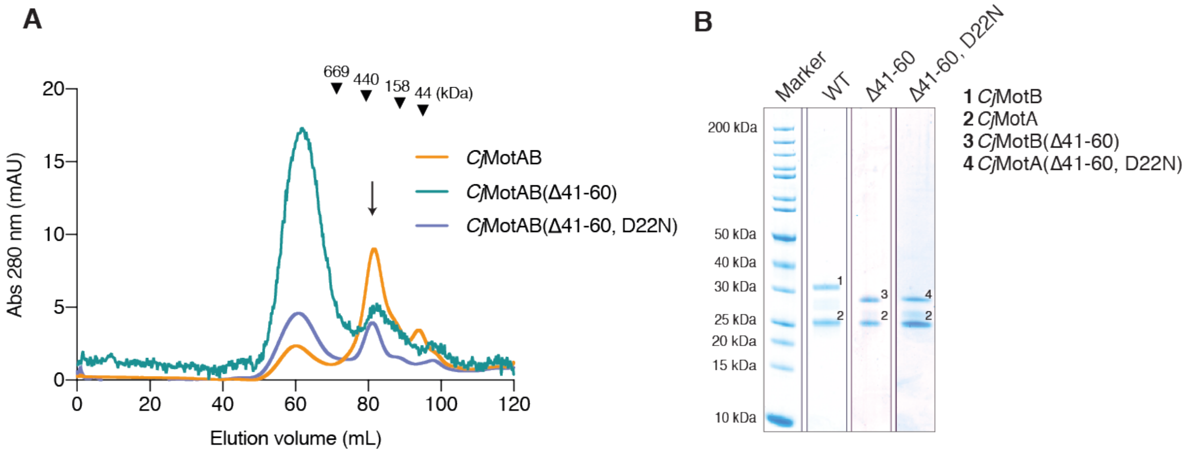

**Fig. S16. Purification of *Cj*MotAB mutants.** (**A**) Size-exclusion chromatography (SEC) profile of detergent-purified *Cj*MotAB, *Cj*MotAB(Δ41-60) and *Cj*MotAB(Δ41-60, D22N) complexes. The fraction used for cryo-EM grid preparation is indicated with an arrow. Elution volumes of molecular weight standards are indicated with inverted triangles. (**B**) The corresponding SDS-PAGE gels for (A) are shown.

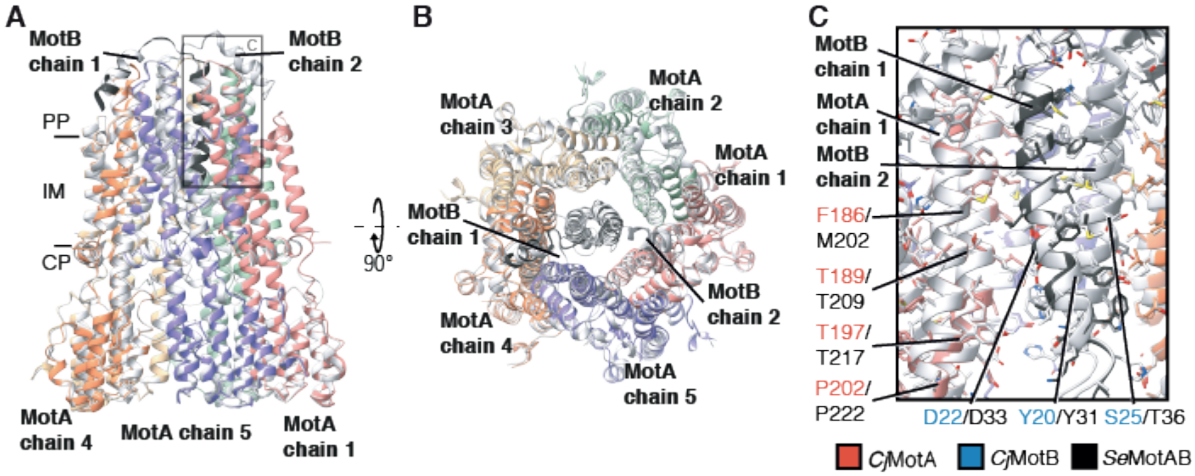

**Fig. S17. Superposition of *Se*MotAB homology model and the *Cj*MotAB cryo-EM structure.** (**A**) Structure of *Se*MotAB, modeled using Modeller based on the *Cj*MotAB cryo-EM structure, colored in the same color code as Fig. 1. The *Cj*MotAB structure is colored in white. Residues *Se*MotA 107-123 and 279-295, corresponding to sequences linking H2 and H3 and a C-terminal extension, respectively, are not shown as they are not present in *Cj*MotA. The calculated $C_\alpha$ r.m.s.d. between both structures is 0.760 Å. (**B**) Same as (A) but top view (periplasmic side). (**C**) close up of the squared region in (A) of the MotA-MotB interface of both stator units displaying the high conservation of the structure.

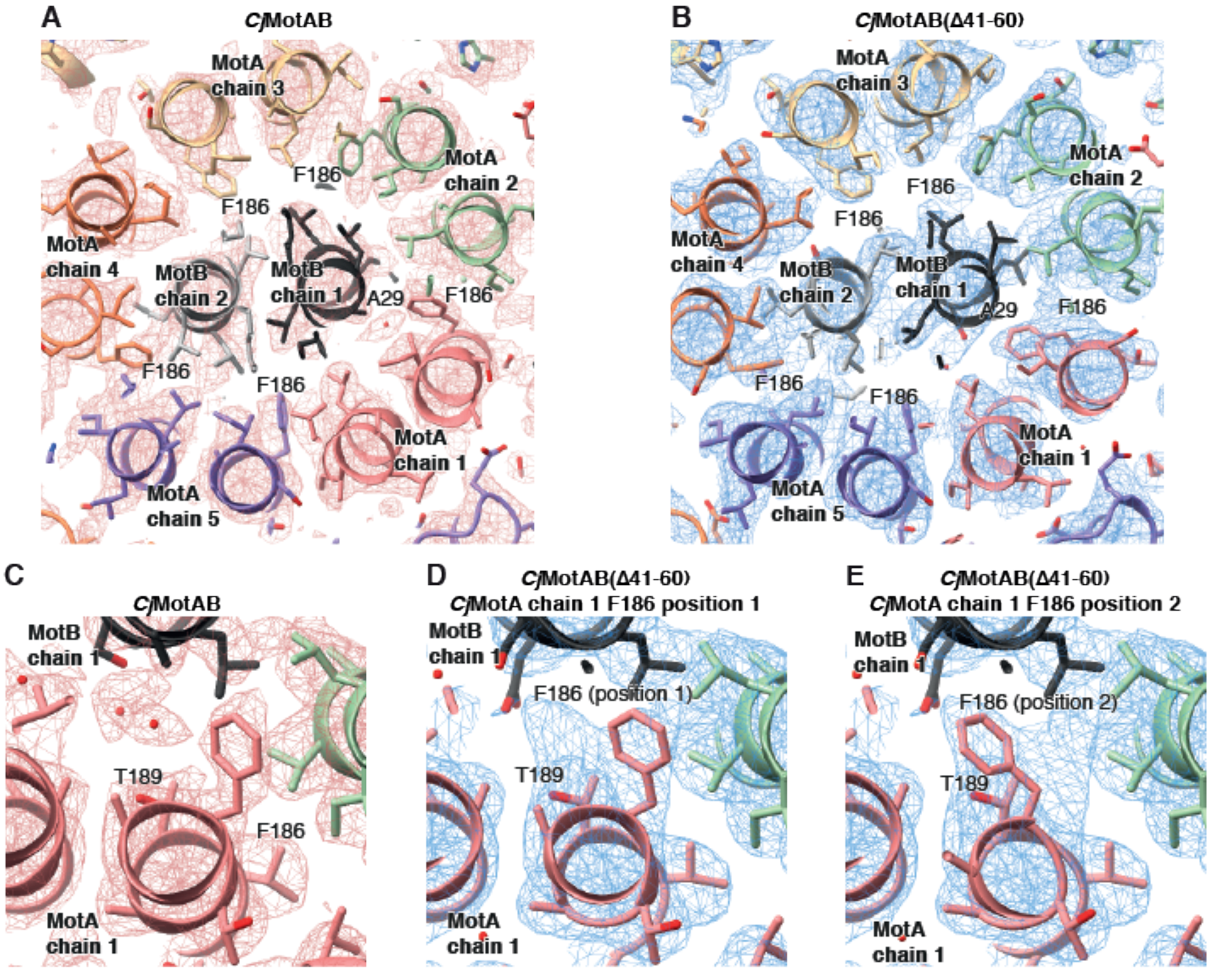

**Fig. S18. Comparison of *Cj*MotA (chain 1) F186 positions in the plugged and unplugged *Cj*MotAB structures.** (**A**) Central view of the (plugged) *Cj*MotAB structure, showing the location and density for all *Cj*MotA F186 residues. (**B**) Same as (A) but showing the (unplugged) *Cj*MotAB(Δ41-60) structure. **(C**) Close-up view of *Cj*MotA chain 1 F186 in the plugged) *Cj*MotAB structure in its only position. (**D**) Close-up view of *Cj*MotA chain 1 F186 in the (unplugged) *Cj*MotAB(Δ41-60) structure in position 1 (same position as in the plugged structure). (**E**) Close-up view of *Cj*MotA chain 1 F186 in the (unplugged) *Cj*MotAB(Δ41-60) structure in position 2. Note that there are also some conformational differences in the main chain around *Cj*MotA chain 1 F186.

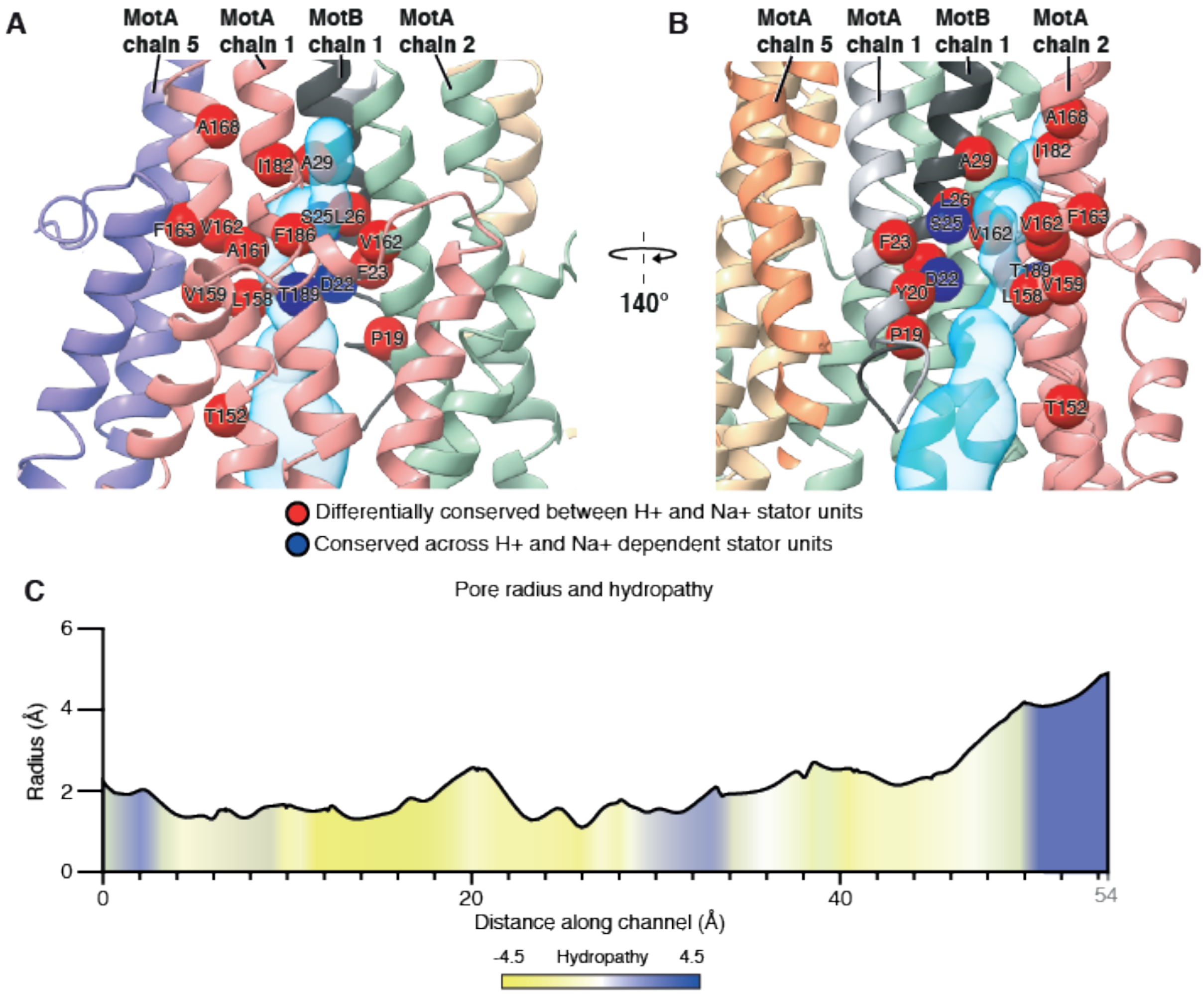

**Fig. S19. The proposed stator unit ion channel.** (**A** and **B**) *Cj*MotAB stator unit (same color code as Fig. 1) with the solvent-accessible volume of the ion channel (cyan) calculated with Mole 2.5 (*63*) (see Methods). Red spheres represent residues that are divergent between $H^+$- and $Na^+$-dependent stator units but that are conserved inside each group, while blue spheres represent highly conserved residues across both $H^+$- and $Na^+$-dependent stator units. (**C**) Profile of channel radius and hydropathy for the calculated ion channel. The channel length is 54 Å and the bottleneck radius is 1.1 Å.

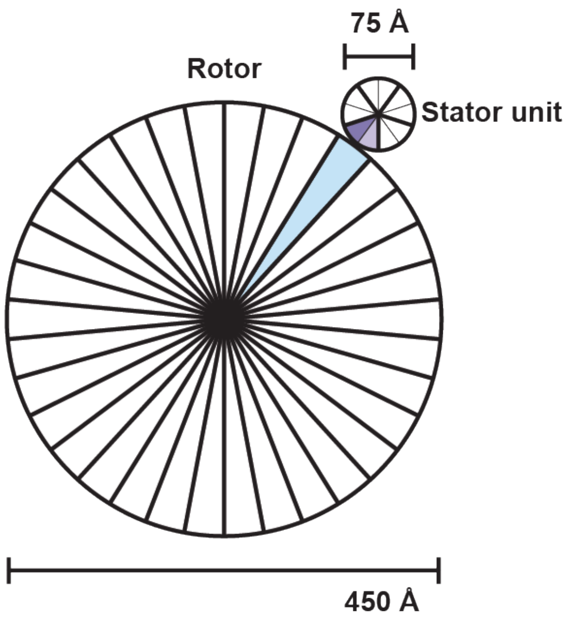

**Fig. S20. Schematic representation of the rotor–stator unit scale, symmetry and interaction.** Schematic representation of the top of the rotor and the stator. The rotor has been shown as a 450 Å disc representing the measured distance (by the authors of the present study) at the top of the C-ring (expected to be the location of FliG $Helix_{Torque}$) of the wild type rotor (*88*). The expected FliG stoichiometry (34-fold) is represented by splitting up the disc in 34 equally-sized slices, one of which is colored in light blue. The stator unit is represented as a disc of diameter 75 Å, the measured diameter of the stator unit at its cytoplasmic region. The 5-fold stoichiometry and pseudo-symmetry of MotA is represented by its division into 5 equally-sized slices with thick black lines. The slices are subdivided into two to represent the proposed movement upon proton or hydronium transport in 36° steps as discussed in the text and in fig. S21. One of the five large slices of the stator unit is colored: one sub-slice is colored in purple, the other in light purple.

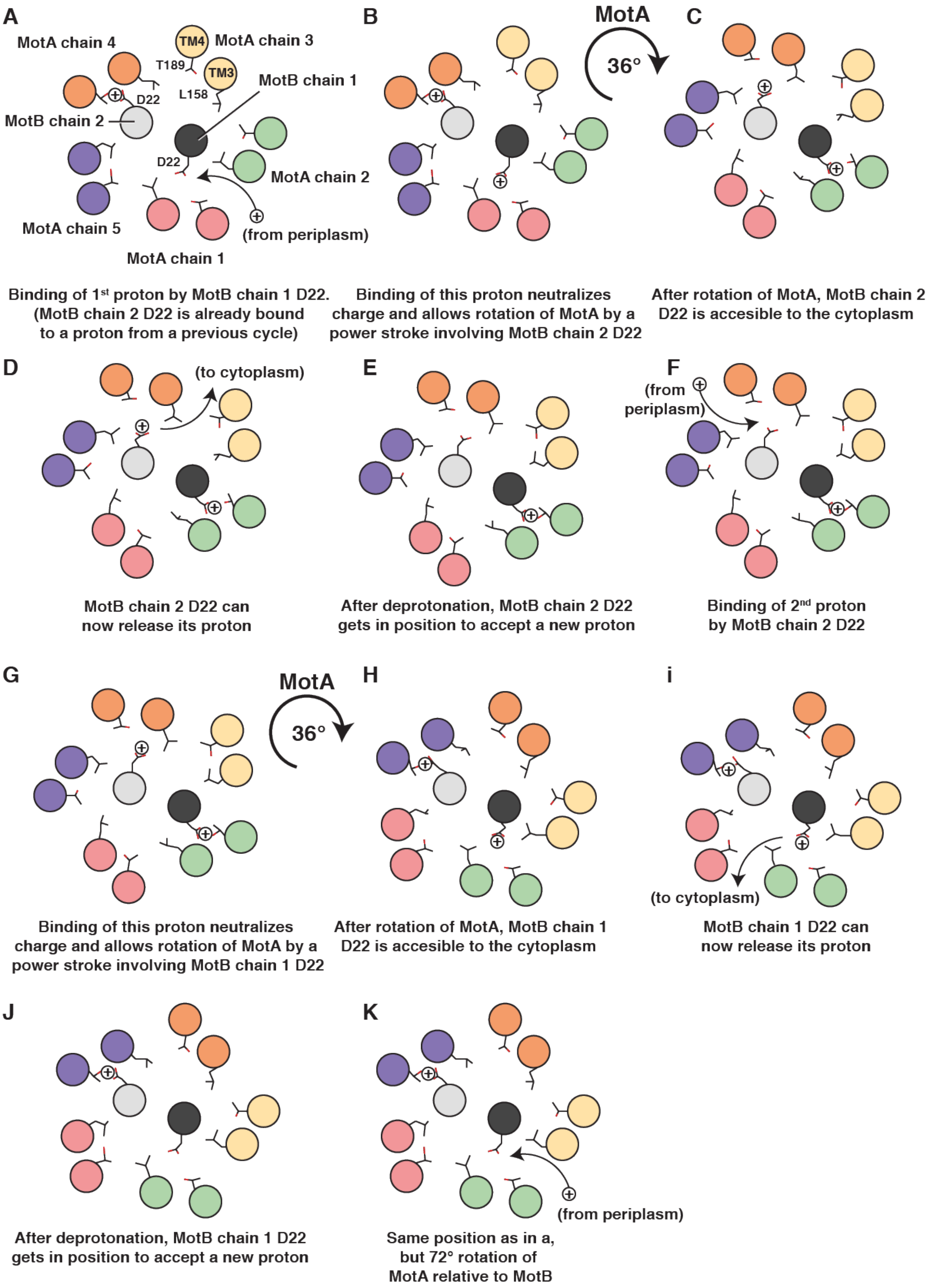
A
MotA chain 4
TM4
MotA chain 3
T189
TM3
MotB chain 1
L158
D22
MotB chain 2
D22
MotA chain 2
MotA chain 5
(from periplasm)
MotA chain 1
Binding of 1st proton by MotB chain 1 D22. (MotB chain 2 D22 is already bound to a proton from a previous cycle)
B
MotA
36°
Binding of this proton neutralizes charge and allows rotation of MotA by a power stroke involving MotB chain 2 D22
C
After rotation of MotA, MotB chain 2 D22 is accesible to the cytoplasm
D
(to cytoplasm)
MotB chain 2 D22 can now release its proton
E
After deprotonation, MotB chain 2 D22 gets in position to accept a new proton
F
(from periplasm)
Binding of 2nd proton by MotB chain 2 D22
G
MotA
36°
Binding of this proton neutralizes charge and allows rotation of MotA by a power stroke involving MotB chain 1 D22
H
After rotation of MotA, MotB chain 1 D22 is accesible to the cytoplasm
i
(to cytoplasm)
MotB chain 1 D22 can now release its proton
J
After deprotonation, MotB chain 1 D22 gets in position to accept a new proton
K
(from periplasm)
Same position as in a, but 72° rotation of MotA relative to MotB

**Fig. S21. Mechanistic model for proton motive force-powered rotation of MotA around MotB.** (**A**) MotB chain 1 D22 binds a proton (or hydronium, represented by a sphere with a + symbol) from the periplasmic side. (**B**) Proton (or hydronium) binding neutralizes charge and MotA rotates 36° clockwise (CW) by a power stroke involving MotB chain 2 D22. (**C**) Now, MotB chain 2 D22 is accessible to the cytoplasm and releases its proton (**D**). (**E**) After deprotonation, MotB chain 2 D22 gets in position and binds a new proton (**F**). (**G**) As before, binding of this proton neutralizes charge, which allows 36° rotation of MotA by a power stroke involving MotB chain 1 D22. (**H**) MotB chain 1 D22 is accessible to the cytoplasm and can now release its proton (**I**). (**J**) After deprotonation, MotB chain 1 D22 gets in position to accept a new proton. (**K**) Same as in (A), but after a 72° rotation of MotA relative to MotB.

#### Tables

#### Table S1. Cryo-EM data collection, refinement and validation statistics

| | *Cj*MotAB | *Cj*MotAB (Δ41-60) | *Cj*MotAB (Δ41-60, D22N) | *So*MotAB | *Va*PomAB |
|---|---|---|---|---|---|
| **Data collection and processing** | | | | | |
| Microscope | | | Titan Krios G2 | | |
| Voltage (kV) | | | 300 | | |
| Magnification (nominal) | | | 96,000x | | |
| Total exposure (e-/Å$^2$) | 40.84 | 42.51 | 43.44 | 40.84 | 42.82 |
| Pixel size (Å) | | | 0.832 | | |
| Defocus range (µm) | | | -1 to -3 | | |
| Micrographs used (no.) | 5,434 | 5,939 | 3,286 | 9,040 | 6,388 |
| Total picked particles (no.) | 2,695,672 | - | 2,043,384 | 5,431,609 | 1,037,574 |
| Final particles (no.) | 278,762 | 329,503 | 456,384 | 799,093 | 91,076 |
| Box size (pixels) | | | 256 | | |
| Symmetry imposed | | | C1 | | |
| Map-sharpening B-factor (Å$^2$) | -117 | -132 | -115 | -115 | -772 |
| Map resolution (Å) (FSC 0.143) | 3.10 | 2.98 | 3.00 | 3.48 | 6.45 |
| **Refinement** | | | | | |
| Model composition | | | | | |
| Non-hydrogen atoms | 10,434 | 10,190 | 10,189 | | |
| Protein residues | 1,357 | 1,326 | 1,326 | | |
| Solvent molecules | 8 | 7 | 6 | | |
| B-factors (Å$^2$) | | | | | |
| Protein | 41.93 | 74.45 | 39.08 | | |
| Solvent | 40.81 | 72.63 | 42.84 | | |
| R.m.s. deviations | | | | | |
| Bond lengths (Å) | 0.009 | 0.014 | 0.005 | | |
| Bond angles (°) | 1.376 | 1.530 | 1.207 | | |
| CC (mask) | 0.86 | 0.85 | 0.84 | | |
| Refinement resolution (FSC map vs. model (masked)=0.143) (Å) | 3.0 | 2.9 | 2.89 | | |
| **Validation** | | | | | |
| MolProbity score | 1.37 | 1.24 | 1.12 | | |
| Clashscore | 4.12 | 2.72 | 2.28 | | |
| Poor rotamers (%) | 0.00 | 0.09 | 0.00 | | |
| Ramachandran plot | | | | | |
| Favored (%) | 97.02 | 96.95 | 97.41 | | |
| Allowed (%) | 2.98 | 2.97 | 2.59 | | |
| Disallowed (%) | 0.00 | 0.08 | 0.00 | | |

**Table S2. Growth of *Salmonella enterica* MotAB point mutants.**

| | Growth rate | Doubling time |
|---|---|---|
| WT | 0.0188 ± 0.0009 min$^{-1}$ | 36.98 ± 1.84 min |
| Δ*motAB* | 0.0180 ± 0.0022 min$^{-1}$ | 39.03 ± 5.26 min |
| ***Se*MotA point mutants** | | |
| MotA R90K | 0.0183 ± 0.0007 min$^{-1}$ | 37.99 ± 1.50 min |
| MotA R90E | 0.0172 ± 0.0013 min$^{-1}$ | 40.47 ± 3.29 min |
| MotA R90A | 0.0185 ± 0.0012 min$^{-1}$ | 37.49 ± 2.45 min |
| MotA R90D | 0.0192 ± 0.0011 min$^{-1}$ | 36.17 ± 2.06 min |
| MotA Q91K | 0.0169 ± 0.0018 min$^{-1}$ | 41.42 ± 4.12 min |
| MotA Q91E | 0.0169 ± 0.0014 min$^{-1}$ | 41.10 ± 3.61 min |
| MotA Q91A | 0.0166 ± 0.0016 min$^{-1}$ | 42.09 ± 3.93 min |
| MotA Q91D | 0.0173 ± 0.0025 min$^{-1}$ | 40.60 ± 6.16 min |
| MotA F95A | 0.0207 ± 0.0010 min$^{-1}$ | 33.52 ± 1.72 min |
| MotA E98D | 0.0181 ± 0.0023 min$^{-1}$ | 38.72 ± 5.31 min |
| MotA E98R | 0.0179 ± 0.0012 min$^{-1}$ | 38.74 ± 2.53 min |
| MotA-E98K | 0.0179 ± 0.0032 min$^{-1}$ | 39.45 ± 6.40 min |
| MotA E98A | 0.0168 ± 0.0012 min$^{-1}$ | 41.44 ± 2.79 min |
| MotA S135R | 0.0201 ± 0.0015 min$^{-1}$ | 34.53 ± 2.39 min |
| MotA S135A | 0.0178 ± 0.0024 min$^{-1}$ | 39.36 ± 4.93 min |
| MotA S135E | 0.0176 ± 0.0019 min$^{-1}$ | 39.69 ± 4.25 min |
| MotA E155R | 0.0183 ± 0.0024 min$^{-1}$ | 38.31 ± 5.40 min |
| MotA E155A | 0.0173 ± 0.0004 min$^{-1}$ | 40.19 ± 0.85 min |
| MotA E155Q | 0.0169 ± 0.0012 min$^{-1}$ | 41.16 ± 2.93 min |
| MotA E155D | 0.0159 ± 0.0017 min$^{-1}$ | 43.77 ± 4.40 min |
| MotA P173A | 0.0154 ± 0.0009 min$^{-1}$ | 45.19 ± 2.48 min |
| MotA A191E | 0.0170 ± 0.0020 min$^{-1}$ | 41.04 ± 4.53 min |
| MotA A191R | 0.0187 ± 0.0010 min$^{-1}$ | 37.10 ± 2.01 min |
| MotA P194A | 0.0191 ± 0.0015 min$^{-1}$ | 36.40 ± 2.82 min |
| MotA M206A | 0.0171 ± 0.0013 min$^{-1}$ | 40.61 ± 3.29 min |
| MotA M206D | 0.0163 ± 0.0006 min$^{-1}$ | 42.63 ± 1.61 min |
| MotA T209A | 0.0181 ± 0.0011 min$^{-1}$ | 38.48 ± 2.48 min |
| MotA T209S | 0.0166 ± 0.0013 min$^{-1}$ | 41.85 ± 3.43 min |
| MotA Y217N | 0.0174 ± 0.0016 min$^{-1}$ | 40.08 ± 3.53 min |
| MotA Y217A | 0.0164 ± 0.0011 min$^{-1}$ | 42.42 ± 3.03 min |
| MotA Y217F | 0.0173 ± 0.0008 min$^{-1}$ | 40.10 ± 1.76 min |
| MotA P222A | 0.0164 ± 0.0009 min$^{-1}$ | 42.42 ± 2.19 min |
| MotA P254A | 0.0180 ± 0.0015 min$^{-1}$ | 38.67 ± 3.22 min |
| MotA P255E | 0.0183 ± 0.0022 min$^{-1}$ | 38.21 ± 4.91 min |
| MotA P255A | 0.0173 ± 0.0022 min$^{-1}$ | 40.51 ± 5.59 min |
| MotA L276A | 0.0183 ± 0.0029 min$^{-1}$ | 38.65 ± 6.74 min |
| MotA E294A | 0.0178 ± 0.0005 min$^{-1}$ | 38.86 ± 1.01 min |
| MotA E294K | 0.0191 ± 0.0030 min$^{-1}$ | 37.01 ± 6.32 min |
| ***Se*MotB point mutants** | | |
| MotB R13A | 0.0177 ± 0.0017 min$^{-1}$ | 39.30 ± 3.75 min |
| MotB R13E | 0.0182 ± 0.0022 min$^{-1}$ | 38.36 ± 4.41 min |
| MotB W27A | 0.0175 ± 0.0015 min$^{-1}$ | 39.80 ± 3.35 min |
| MotB Y31A | 0.0190 ± 0.0018 min$^{-1}$ | 36.76 ± 3.72 min |
| MotB Y31F | 0.0186 ± 0.0021 min$^{-1}$ | 37.51 ± 4.05 min |
| MotB D33A | 0.0189 ± 0.0015 min$^{-1}$ | 36.86 ± 3.08 min |
| MotB D33E | 0.0169 ± 0.0010 min$^{-1}$ | 41.16 ± 2.54 min |
| MotB D33N | 0.0168 ± 0.0016 min$^{-1}$ | 41.46 ± 3.77 min |
| MotB T36A | 0.0181 ± 0.0019 min$^{-1}$ | 38.67 ± 4.00 min |
| MotB S49A | 0.0178 ± 0.0027 min$^{-1}$ | 39.52 ± 6.05 min |
| MotB E55A | 0.0186 ± 0.0019 min$^{-1}$ | 37.61 ± 3.92 min |
| MotB E55K | 0.0181 ± 0.0008 min$^{-1}$ | 38.34 ± 1.68 min |
| MotB L56E | 0.0183 ± 0.0014 min$^{-1}$ | 37.97 ± 2.86 min |
| MotB L56C | 0.0190 ± 0.0014 min$^{-1}$ | 36.58 ± 2.56 min |
| MotB I59E | 0.0188 ± 0.0009 min$^{-1}$ | 36.91 ± 1.90 min |
| MotB I59C | 0.0193 ± 0.0018 min$^{-1}$ | 36.06 ± 3.32 min |

| | | | | | | |
|---|---|---|---|---|---|---|
| MotB F63R | 0.0199 | ± 0.0014 | $min^{-1}$ | 34.95 | ± 2.34 | min |
| MotB F63C | 0.0210 | ± 0.0008 | $min^{-1}$ | 33.03 | ± 1.27 | min |

WT, wildtype; Δ*motAB*, *motAB* clean deletion; *Se*MotAB, *Salmonella enterica* MotAB.

The growth rate data represents the mean values ± standard deviation of three biological replicates.

**References**

64. P. Ma *et al.*, An efficient strategy for small-scale screening and production of archaeal membrane transport proteins in Escherichia coli. *PloS one* **8**, e76913 (2013).

65. J. Zivanov *et al.*, New tools for automated high-resolution cryo-EM structure determination in RELION-3. *eLife* **7**, 163 (2018).

66. S. Q. Zheng *et al.*, MotionCor2: anisotropic correction of beam-induced motion for improved cryo-electron microscopy. *Nature methods* **14**, 331-332 (2017).

67. A. Rohou, N. Grigorieff, CTFFIND4: Fast and accurate defocus estimation from electron micrographs. *Journal of structural biology* **192**, 216-221 (2015).

68. A. Punjani, J. L. Rubinstein, D. J. Fleet, M. A. Brubaker, cryoSPARC: algorithms for rapid unsupervised cryo-EM structure determination. *Nature methods* **14**, 290-296 (2017).

69. T. Grant, A. Rohou, N. Grigorieff, cisTEM, user-friendly software for single-particle image processing. *Elife* **7**, (2018).

70. Y. Z. Tan *et al.*, Addressing preferred specimen orientation in single-particle cryo-EM through tilting. *Nature methods* **14**, 793-796 (2017).

71. P. Emsley, B. Lohkamp, W. G. Scott, K. Cowtan, Features and development of Coot. *Acta crystallographica Section D, Biological crystallography* **66**, 486-501 (2010).

72. D. Liebschner *et al.*, Macromolecular structure determination using X-rays, neutrons and electrons: recent developments in Phenix. *Acta Crystallographica Section D Structural Biology* **75**, 861-877 (2019).

73. C. J. Williams *et al.*, MolProbity: More and better reference data for improved all-atom structure validation. *Protein science : a publication of the Protein Society* **27**, 293-315 (2018).

74. T. F. Braun, L. Q. Al-Mawsawi, a. Seiji Kojima, D. F. Blair, Arrangement of Core Membrane Segments in the MotA/MotB Proton-Channel Complex of Escherichia coli†. *Biochemistry* **43**, 35-45 (2003).

75. D. F. Blair, D. Y. Kim, H. C. Berg, Mutant MotB proteins in Escherichia coli. *Journal of bacteriology* **173**, 4049-4055 (1991).

76. A. Sali, T. L. Blundell, Comparative protein modelling by satisfaction of spatial restraints. *Journal of molecular biology* **234**, 779-815 (1993).

77. K.-H. Lam *et al.*, Multiple conformations of the FliG C-terminal domain provide insight into flagellar motor switching. *Structure (London, England : 1993)* **20**, 315-325 (2012).

78. B. Raveh, N. London, O. Schueler-Furman, Sub-angstrom modeling of complexes between flexible peptides and globular proteins. *Proteins* **78**, 2029-2040 (2010).

79. N. London, B. Raveh, E. Cohen, G. Fathi, O. Schueler-Furman, Rosetta FlexPepDock web server--high resolution modeling of peptide-protein interactions. *Nucleic acids research* **39**, W249-253 (2011).

80. J. E. Karlinsey, lambda-Red genetic engineering in Salmonella enterica serovar Typhimurium. *Methods in enzymology* **421**, 199-209 (2007).

81. K. E. Sanderson, J. R. Roth, Linkage map of Salmonella typhimurium, edition VII. *Microbiological reviews* **52**, 485-532 (1988).

82. C. A. Schneider, W. S. Rasband, K. W. Eliceiri, NIH Image to ImageJ: 25 years of image analysis. *Nature methods* **9**, 671-675 (2012).

83. B. G. Hall, H. Acar, A. Nandipati, M. Barlow, Growth rates made easy. *Molecular Biology and Evolution* **31**, 232-238 (2014).

84. E. F. Pettersen *et al.*, UCSF Chimera--A visualization system for exploratory research and analysis. *Journal of Computational Chemistry* **25**, 1605-1612 (2004).

85. T. D. Goddard *et al.*, UCSF ChimeraX: Meeting modern challenges in visualization and analysis. *Protein science : a publication of the Protein Society* **27**, 14-25 (2018).

86. E. Jurrus *et al.*, Improvements to the APBS biomolecular solvation software suite. *Protein science : a publication of the Protein Society* **27**, 112-128 (2018).

87. H. Ashkenazy *et al.*, ConSurf 2016: an improved methodology to estimate and visualize evolutionary conservation in macromolecules. *Nucleic acids research* **44**, W344-350 (2016).

88. T. Sakai et al., Novel Insights into Conformational Rearrangements of the Bacterial Flagellar Switch Complex. mBio 10, 19 (2019).

**Movie S1. Structural organization of the flagellar stator unit *Cj*MotAB.** The movie begins showing the cryo-EM density map for the *Cj*MotAB stator unit in the micelle (grey), followed by color representation of the same map with and without the micelle, rotating around the vertical axis for visualization purposes. Finally, the atomic model is shown highlighting the 5:2 stoichiometry of the complex by rotating 90° around the horizontal axis to show first top (periplasmic side) and then bottom (cytoplasmic side) views. Color code is the same as in Fig. 1.

**Movie S2. (Un)plugging of the flagellar stator unit.** Conformational changes of the atomic model of *Cj*MotAB along the process of (un)plugging. First, the plugged structure is presented, and the plugs are hidden for the next steps. This is followed by the visualization of the conformational changes upon unplugging and plugging on the overall structure of the stator unit. The view is then focused to show the conformational changes upon plugging and unplugging on the interface between MotA chain 1 and chain 2, showing residue F186 of MotA chain 1 (visualized in the unplugged structure in its position 2). Subsequently, with the complex in unplugged conformation, the flexibility of MotA chain1 F186 is shown by morphing the structures between position 1 and 2. The view is then rotated 180° around the vertical axis to visualize the F186 flexibility from the inside of the complex. Finally, the channel proposed on this study is shown, first by following its length along the inside of the complex and finally providing the whole view of it, showing the residues lining the channel. Color code is the same as in Fig. 1.

**Movie S3. Effect of (mimicking) proton or hydronium binding to the flagellar stator unit.** Conformational changes of the atomic model of *Cj*MotAB between deprotonated and protonated states. The deprotonated state is the *Cj*MotAB(Δ41-60) structure. The protonated state shown is the *Cj*MotAB(Δ41-60, N22) structure, but displaying MotB N22 as aspartate. The movie begins with the general view of the stator unit in its unplugged, deprotonated state. The structure then morphs back and forth between the deprotonated and protonated states (this is not expected to occur in reality, we use this movement for visualization purposes only). The deprotonated stator unit is then turned and focused on the MotB dimer. It can be appreciated how the MotB chain 1 D22 changes from pointing to the periplasm on the deprotonated state to pointing to the cytoplasm when protonated by morphing both states between them. The movie ends at the initial position. Color code is the same as in Fig. 1.

**Movie S4. Proposed mechanism of action of the stator unit complex: scheme.** This is an animated version of fig. S21, which shows a model for MotAB mechanism of action where rotation of MotA is caused by a power stroke upon protonation of MotB D22. First, MotB chain 1 D22 binds a proton (or hydronium, represented by a sphere with a + symbol) from the periplasmic side. Proton (or hydronium) binding neutralizes charge, which allows 36° rotation of MotA clockwise (CW) relative to MotB, by a power stroke involving MotB chain 2 D22. After rotation of MotA, MotB chain 2 D22 is accessible to the cytoplasm and releases its proton. After deprotonation, MotB chain 2 D22 gets in position and binds a new proton. As before, binding of this proton neutralizes charge, which allows another 36° rotation of MotA by a power stroke involving MotB chain 1 D22. MotB chain 1 D22 is now accessible to the cytoplasm and can release its proton. After deprotonation, MotB chain 1 D22 gets in position to accept a new proton, which is the same situation as in the beginning, but after a 72° rotation of MotA relative to MotB. In the movie, this process repeats until the stator unit is back in the starting position, after transport of 10 protons. Color code is the same as in Fig. 1.

**Movie S5. Proposed mechanism of action of the stator unit complex: modeling.** MotAB stator unit is first shown from the side view, from which it is rotated 90° around the horizontal axis to show the periplasmic view of the complex. Then, the model for the 36°-step turning of MotA pentamer around the MotB dimer is shown. Protonation of MotB chain 1 D22 (note that MotB chain 2 D22 is already protonated from a previous cycle) allows the power stroke produced by MotB chain 2, which gives place to a 36° rotation of the MotA pentamer. Consecutively, MotB chain 2 D22 is protonated, allowing the power stroke performed by MotB chain 1. The same process is shown in more detail with the slice of the complex at the level of the D22 residues of both MotB chains. MotB chain 2 D22 is engaged at MotA chain 2 (represented by a cyan halo). The incoming proton (or hydronium, represented by a sphere with a + symbol) from the cytoplasm, binds to MotB chain 1 D22, neutralizing the charge. This allows the 36° rotation of the MotA pentamer around MotB, given by the power stroke produced by MotB chain 2. Now, MotB chain 1 is engaged at MotA chain 5. At the same time, the proton interacting with MotB chain 2 is released to the cytoplasm. The model is followed by a change of roles between MotB chain 1 and 2. The same process is shown from the side, first as the whole complex and later as a sliced view highlighting the role of the D22 residue of both MotB chains. Finally, the view ends on the initial position. Color code is the same as in Fig. 1.
